# A descending inhibitory mechanism of nociception mediated by an evolutionarily conserved neuropeptide system in *Drosophila*

**DOI:** 10.1101/2022.03.08.483420

**Authors:** Izumi Oikawa, Shu Kondo, Kao Hashimoto, Akiho Yoshida, Megumi Hamajima, Hiromu Tanimoto, Katsuo Furukubo-Tokunaga, Ken Honjo

## Abstract

Nociception is a neural process that animals have developed to avoid potentially tissue-damaging stimuli. While nociception is triggered in the peripheral nervous system, its modulation by the central nervous system is a critical process in mammals, whose dysfunction has been extensively implicated in chronic pain pathogenesis. The peripheral mechanisms of nociception are largely conserved across the animal kingdom. However, it is unclear whether the brain-mediated modulation is also conserved in non-mammalian species. Here, we show that *Drosophila* has a descending inhibitory mechanism of nociception from the brain, mediated by the neuropeptide Drosulfakinin (DSK), a homolog of cholecystokinin (CCK) that plays an important role in the descending control of nociception in mammals. We found that mutants lacking *dsk* or its receptors are hypersensitive to noxious heat. Through a combination of genetic, behavioral, histological, and Ca^2+^ imaging analyses, we subsequently revealed neurons involved in DSK-mediated nociceptive regulation at a single-cell resolution and identified a DSKergic descending neuronal pathway that inhibits nociception. This study provides the first evidence for a descending modulatory mechanism of nociception from the brain in a non-mammalian species that is mediated by the evolutionarily conserved CCK system, raising the possibility that the descending inhibition is an ancient mechanism to regulate nociception.

## Introduction

Minimizing tissue damage is a fundamental task for all animals to increase their chance of survival. Thus, elucidating the principles of nociception, the neural process detecting and encoding potentially tissue-damaging stimuli, is critical to understanding the molecular and neural mechanisms implementing adaptive behaviors and their evolution. Nociceptors are sensory neurons specialized to detect harmful stimuli, whose activation triggers downstream nociceptive circuits and nocifensive responses (1). Since the activities of nociceptors and downstream nociceptive circuits are tightly linked to pain perception in humans, unveiling the mechanisms of nociception is also crucial to a better understanding of human pain mechanisms (2, 3).

Descending inhibition has been suggested to be a pivotal mechanism in the modulation of nociception and pain in mammals. Since the discovery that electrical stimulations of parts of the midbrain in rats enabled surgical operations without anesthetics (4), mammalian descending nociceptive pathways have been implicated in various analgesic phenomena/treatments and the development of chronic pain states, suggesting their critical role in modulating nociception and pain (5, 6). However, brain-mediated modulatory mechanisms of nociception such as descending inhibition have currently been identified only in mammals, despite a high degree of commonality across species in the peripheral nociceptive mechanisms (7, 8). Therefore, it is unknown whether the descending modulation is a *de novo* mechanism typical of the highly developed mammalian central nervous system (CNS) or a conserved control also present in simpler animals.

The descending nociceptive-modulatory systems in mammals have been revealed to involve various neurochemical pathways (5, 6, 9); among these, the cholecystokinin (CCK) system is one of the most extensively characterized (6, 9). In rodents, CCK signaling plays a crucial role in facilitating nociception by counteracting the opioidergic systems in the periaqueductal gray (PAG)—rostral ventral medulla (RVM)—spinal descending pathway (10–13) and inhibiting nociception through the central amygdala (CeA)—PAG—spinal pathway (14). In humans, CCK signaling has been implicated in nocebo hyperalgesia, mediated by the descending nociceptive control system (15). The CCK system is very well-conserved and has been implicated in several common physiological functions among bilaterian species (16–20). However, whether CCK is functionally involved in regulating nociception outside of mammals remains unknown.

Drosulfakinin (DSK), a neuropeptide homologous to CCK, was identified in the fruit fly *Drosophila melanogaster* (19). Fly DSK is reportedly involved in modulating many physiological functions shared with mammals, including gut functions, anxiety, aggression, memory, feeding, synaptic functions, and courtship behaviors (19, 21–23). After the discovery of stereotyped nociceptive escape behavior called rolling and polymodal Class IV md (C4da) nociceptors (24, 25), the larval *Drosophila* has been successfully utilized to identify evolutionarily conserved and previously uncharacterized molecular pathways in nociception (26–34) and circuitry mechanisms in the ventral nerve cord (VNC; the invertebrate equivalent of the spinal cord) to compute multimodal sensory stimuli and select nociceptive escape strategies (35–40). Previous studies have demonstrated that neuropeptidergic systems also participate in regulating nociception in *Drosophila* (39, 41–44). However, the role of fly DSK in nociception remains elusive.

Here, using a collective approach of genetic, behavioral, histological, and Ca^2+^ imaging analyses, we pursued the mechanisms of DSK-mediated nociceptive regulation and demonstrate that the DSK system constitutes a descending inhibitory pathway of nociception from the brain to the VNC in larval *Drosophila*.

## Results

### DSK signaling negatively regulates thermal nociception

Through a thermal nociception screen using the nocifensive rolling response of *Drosophila* larvae, we found that a deletion mutant line of the *dsk* gene showed thermal hypersensitivity with a significantly shorter latency in their response to a 42 °C probe than the controls, suggesting that DSK plays a role in negatively regulating nociception (Figure 1A and B). A genomic fragment containing the wild-type *dsk* gene, and no other neuropeptide genes, significantly rescued the thermal hypersensitivity of the *dsk* mutants (Figure 1B), confirming that *dsk* is responsible for the thermal hypersensitivity.

**Figure 1.**
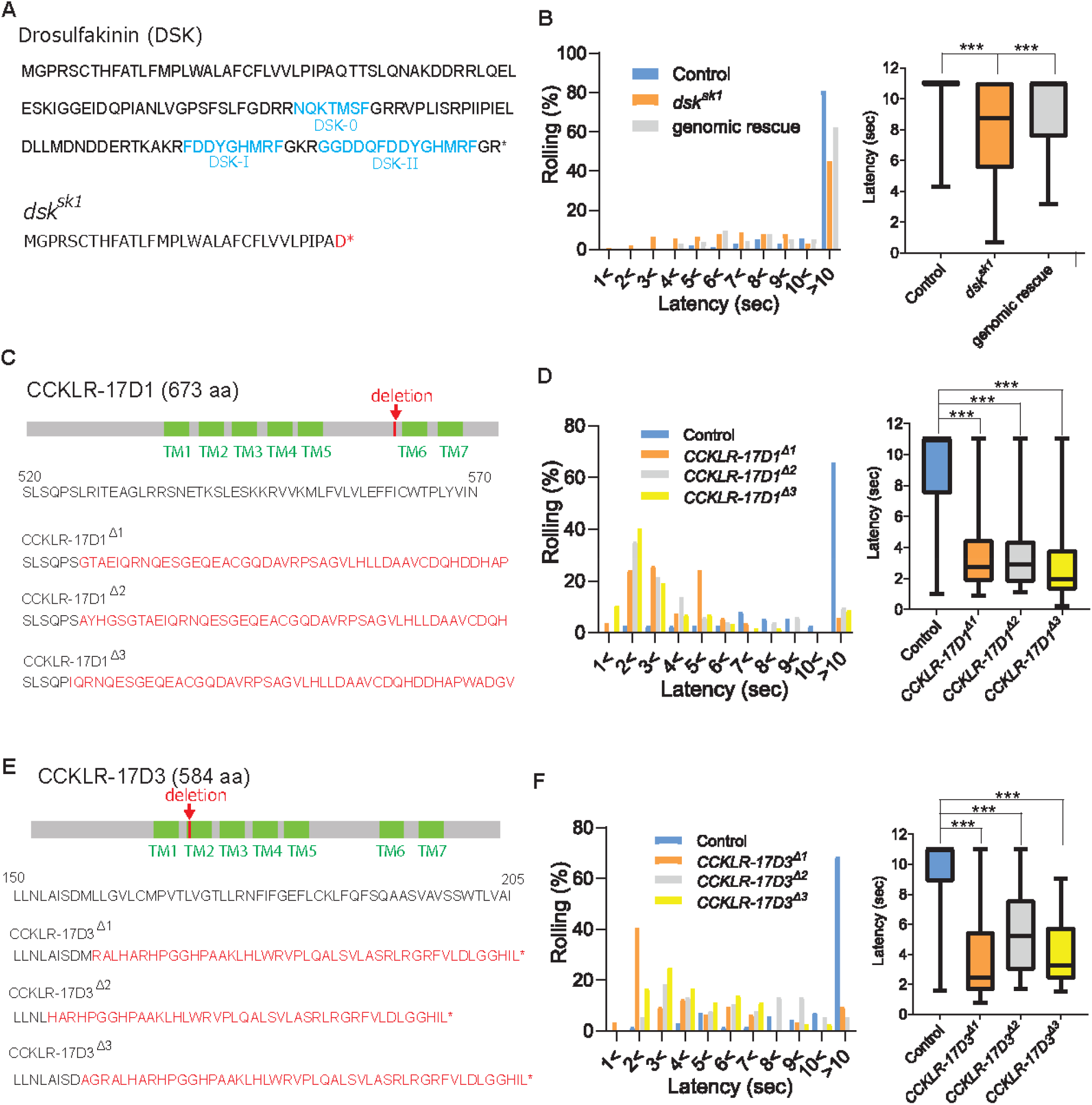
DSK signaling is involved in negatively regulating nociception. (A) Predicted amino acid sequences of pro-DSK peptide in the wild-type (top) and *dsk^sk1^* mutants (bottom). Due to a 5-base deletion at the +94-99 position in the coding sequence, *dsk^sk1^*mutants are predicted to produce a largely truncated pro-DSK peptide unable to be processed to active DSK peptides. Red letters represent residues different from the wild-type and asterisks indicate stop codons. (B) Hypersensitivity of *dsk^sk1^* mutants to a 42°C thermal probe. Significantly shortened latencies of *dsk^sk1^* mutants (n = 143) compared with the controls (n = 164) were recovered in genomic rescue animals (*Dp(3;1)2-2*; *dsk^sk1^*, n = 139). (Left) Histograms (Right) Box plots of latencies. *** p < 0.001 Steel’s test. (C) Predicted amino acid sequence of CCKLR-17D1 in the wild-type and *CCKLR-17D1* mutants. *CCKLR-17D1^Δ1^*, *CCKLR-17D1^Δ2^*, and *CCKLR-17D1^Δ3^* have 17-base (+1573-1589), 2-base (+1575-1576) and 32-base (+1573-1604) deletions in the coding sequence respectively, which result in large frameshifts completely abolishing the sixth and seventh transmembrane domains (TMs). (D) *CCKLR-17D1^Δ1^*(n = 54), *CCKLR-17D1^Δ2^* (n = 51) and *CCKLR-17D1^Δ3^*mutants (n = 57) all showed significantly shorter latencies than control (n = 38). (Left) Histograms (Right) Box plots of latencies. *** p < 0.001 Steel’s test. (E) Predicted amino acid sequence of CCKLR-17D3 in the wild-type and *CCKLR-17D3* mutants. *CCKLR-17D3^Δ1^*, *CCKLR-17D3^Δ2^*, and *CCKLR-17D3^Δ3^* possess 8-base (+474-481), 32-base (+457-488), and 5-base (+472-476) deletions in the coding sequence respectively, which result in large frameshifts and truncation in the middle of the second TM. (F) *CCKLR-17D3^Δ1^* (n = 32), *CCKLR-17D3^Δ2^* (n = 38), and *CCKLR-17D3^Δ3^*mutants (n = 36) all exhibited significantly shorter latencies than control (n = 70). (Left) Histograms (Right) Box plots of latencies. *** p < 0.001 Steel’s test. All box plots show median (middle line) and 25th to 75th percentiles with whiskers indicating the smallest to the largest data points.

DSK has been shown to activate two G-protein coupled receptors, CCKLR-17D1 and CCKLR-17D3 (45, 46), which are orthologous to the mammalian CCK receptors, CCKAR (also known as CCK_1_) and CCKBR (also known as CCK_2_) (16–18). To test whether these receptors mediate DSK signaling in nociception, we generated deletion mutants for *CCKLR-17D1* and *CCKLR-17D3* using CRISPR/Cas9 genome editing and tested them for thermal nociception (Figure 1C-F). When stimulated with a 42 °C probe, three independent deletion lines of the *CCKLR-17D1* and *CCKLR-17D3* exhibited thermal hypersensitivity (Figure 1D and F), further supporting the role of DSK signaling in the negative regulation of thermal nociception. To examine whether the phenotypes of *CCKLR-17D1* and *CCKLR-17D3* mutants are classified as hyperalgesia (hypersensitivity to normally noxious stimuli) or allodynia (abnormal hypersensitivity to normally innocuous stimuli), we tested the receptor mutants with a 38 °C probe, which is close to the threshold of larval thermal nociception (39 °C) (24). We found that the responses of the DSK receptor mutants were indistinguishable from the controls, indicating that the DSK receptor mutants are hypersensitive to suprathreshold thermal stimuli, thus hyperalgesic (Figure 1-figure supplement 1).

We have noticed that the *yw* control strain, which was used by us to generate the *dsk* and receptor deletion mutants, showed relatively longer response latencies to the 42 °C probe compared to the other control strains used in this study. This may be attributed to the effect of the genetic background, although, presently, the cause for this difference is unknown.

### Two groups of brain neurons expressing DSK are responsible for regulating nociception

We attempted to identify DSK-expressing cells in the larval CNS that are responsible for regulating nociception. Unlike mammalian CCK, which is expressed in the CNS and gastrointestinal system, *Drosophila* DSK is expressed in the CNS but not in the gut (19, 47–50). Previous immunohistochemical studies have reported putative DSK-expressing cells in the larval CNS (47, 49). However, since the specificity of the DSK antibodies has not been validated with null mutants in the former studies, it has been a concern that the reported DSK-expressing cells could include non-DSK cells that express the other neuropeptides sharing the C-terminal RFa motif with DSK (51). Consistent with this concern, we found that an antibody against crustacean FLRFa, an FMRFa-like neuropeptide with the C-terminal RFa motif, gives rise to a comparable staining pattern to that of the previously reported DSK antibodies, visualizing cells designated as insulin-producing cells (IPCs; referred to as SP3 in Nichols and Lim, (1996) (47)), MP1, SP1, SP2, Sv, SE2, Tv1-3, T2dm, and A8 in the larval CNS (Figure 2A) (47, 49).

**Figure 2.**
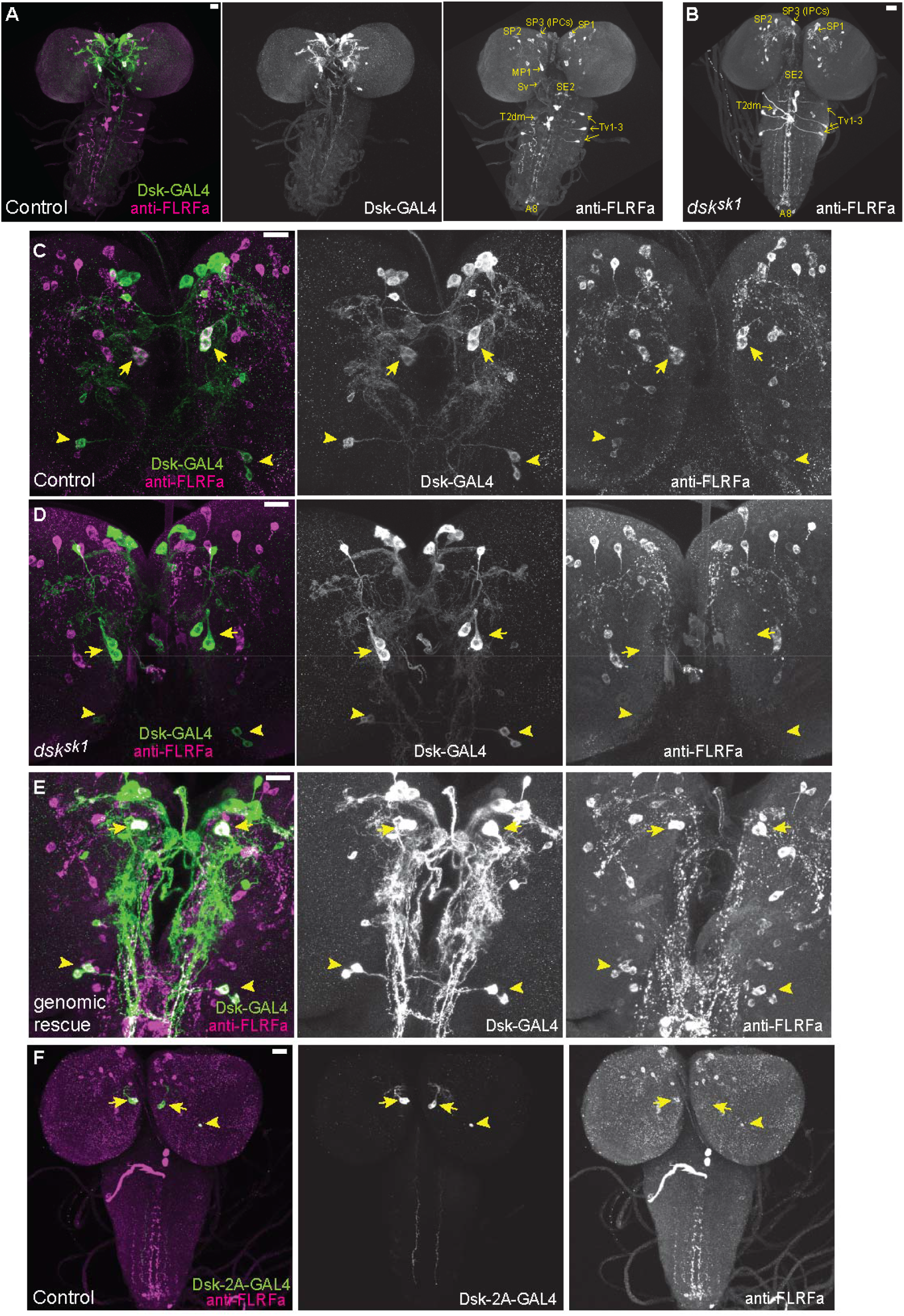
DSK is expressed in two groups of larval brain neurons. (A) Representative image of *DSK-GAL4* expression and anti-FLRFa staining in the wild-type larval CNS. (B) Representative image of anti-FLRFa staining in *dsk^sk1^* larval CNS. (C) Representative images of *DSK-GAL4* expression and anti-FLRFa staining in the control larval brains. Arrows and arrowheads indicate MP1 and Sv neurons, respectively. Co-expressions of DSK-GAL4 and anti-FLRFa were observed in MP1 and Sv in all of the examined samples (n = 27/27), but in IPCs only in 7% (n = 2/27). (D) Representative images of *DSK-GAL4* expression and anti-FLRFa staining in the *dsk^sk1^* larval brains. Arrows and arrowheads indicate MP1 and Sv neurons, respectively. (E) Representative images of *DSK-GAL4* expression and anti-FLRFa staining in the larval brains of the genomic rescue genotype (*Dp(3;1)2-2*/Y; +/*lexA-rCD2::RFP UAS-mCD8::GFP*; *dsk^sk1^ DSK-GAL4*/*dsk^sk1^*). Arrows and arrowheads indicate MP1 and Sv neurons, respectively. (F) An example image showing the expression of *DSK-2A-GAL4*, a 2A-GAL4 knock-in line of the *dsk* gene, more faithfully recapitulating its endogenous expression. Arrows and arrowheads indicate MP1 and Sv neurons, respectively. MP1 neurons were labeled in 100% of *DSK-2AGAL4* samples (n = 41/41), while a single IPC in 51.2% (n = 21/41), multiple IPCs in 24.4% (n = 10/41), and Sv neurons in 17.1% (n = 7/41) of examined samples. All scale bars represent 20 µm.

To identify bona fide DSK-expressing cells responsible for regulating nociception, we performed anti-FLRFa staining in multiple *dsk* null alleles and found that the staining signals persist in all but two pairs of neurons, MP1 and Sv, in the mutant CNS (Figure 2B-D and Figure 2-figure supplement 1A-D), suggesting that these two pairs of neurons are the only neurons in the larval CNS that express DSK. The genomic rescue fragment that rescued the thermal hypersensitivity of *dsk* mutants (Figure 1B) also restored the anti-FLRFa signals in MP1 and Sv in a *dsk* mutant background (Figure 2E), suggesting that the *dsk* gene is responsible for the anti-FLRFa signals in these neurons as well as the thermal nociceptive responses of larvae. This expression pattern was further corroborated by the transgenic reporter line *DSK-GAL4* and the 2A-GAL4 knock-in reporter line *DSK-2A-GAL4* (52), both of which we found were expressed only in MP1 and Sv neurons among the anti-FLRFa-positive neurons (Figure 2A, C, and F; Figure 2-figure supplement 1B-F). We also found that anti-FLRFa, *DSK-GAL4*, and *DSK-2A-GAL4* do not visualize larval peripheral neurons including C4da nociceptors (Figure 2A and F; Figure 2-figure supplement 1G). Taken together, these results identified two sets of brain neurons, MP1 and Sv, as the DSK-expressing cells that are involved in regulating nociception in larvae.

### CCKLR-17D1 in Goro neurons functions to negatively regulate nociception

We sought the potential target cells of DSK signaling for regulating nociception. Since DSK receptor mutants were thermally hypersensitive consistently to *dsk* mutants (Figure 1B, D, and F), neurons in the larval nociceptive circuit that express DSK receptors were promising candidates. To visualize the cells expressing DSK receptors, we generated T2A-GAL4 knock-ins in *CCKLR* genes. Both *CCKLR-17D1-T2A-GAL4* and *CCKLR-17D3-T2A-GAL4* were widely expressed in the larval CNS, predominantly in neuronal cells (Figure 3A and B; Figure 3-figure supplement 1A-D). By performing double-labeling experiments, we found that *CCKLR-17D1-T2A-GAL4* and *CCKLR-17D3-T2A-GAL4* are not expressed in the nociceptors at the periphery (Figure 3C and D; Figure 3-figure supplement 1E and F), or in nociceptive interneurons including Basin1-4 (53), A08n (36, 41), and DnB neurons (35) in the larval VNC (Figure 3-figure supplement 1G-L). However, we found that they are expressed in Goro neurons (Figure 3E and F), which are the fourth-order nociceptive interneurons located in the larval VNC (53).

**Figure 3.**
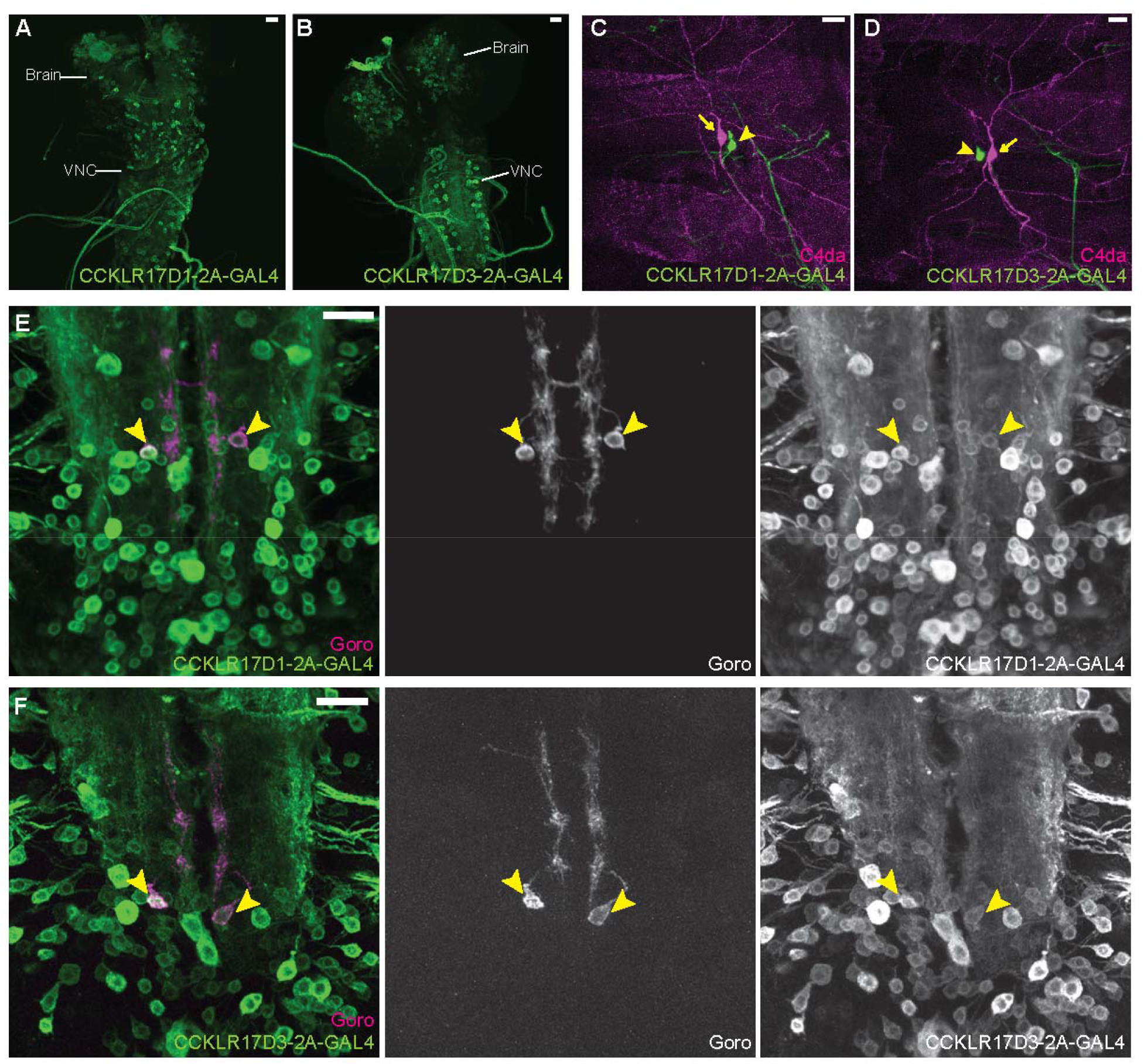
DSK receptors are expressed in Goro neurons in the larval VNC. (A and B) Representative images of *CCKLR-17D1-T2A-GAL4* (A) and *CCKLR-17D3-T2A-GAL4* (B) expressions in the larval CNS. (C and D) Representative images showing double-labeling of *CCKLR-17D1-T2A-GAL4* (C) and *CCKLR-17D3-T2A-GAL4* (D) with C4da nociceptors (*R38A10-lexA*). Arrows and arrowheads indicate cell bodies of C4da nociceptors and es cells, respectively. (E and F) Representative images showing double-labeling of *CCKLR-17D1-T2A-GAL4* (E) and *CCKLR-17D3-T2A-GAL4* (F) with Goro neurons (arrowheads, *R69E06-lexA*). Expression patterns were confirmed in multiple samples (n = 7 and 5). All scale bars represent 20 µm.

To test whether the DSK receptors in Goro neurons are functionally important for regulating nociception, we performed RNAi and rescue experiments using *R69E06-GAL4* that marks Goro neurons (53). RNAi knockdown of CCKLR-17D1, but not CCKLR-17D3, in *R69E06-GAL4* neurons induced thermal hypersensitivity (Figure 4A and Figure 4-figure supplement 1A). *R69E06-GAL4* is also expressed in multiple neurons in the larval brain other than Goro neurons in the VNC (53). However, the thermal hypersensitivity was not observed when the expression of CCKLR-17D1 RNAi was excluded from Goro neurons by *tsh-GAL80*, pointing to the requirement of CCKLR-17D1 in Goro neurons (Figure 4-figure supplement 1B-D). Consistent with these RNAi results, expressing wild-type CCKLR-17D1, but not CCKLR-17D3, with *R69E06-GAL4* rescued the hypersensitivity of the respective mutants (Figure 4B and C). Furthermore, the morphology of Goro neurons was not affected by CCKLR-17D1 knockdown (Figure 4-figure supplement 1E). Therefore, we conclude that CCKLR-17D1 functions in Goro neurons to negatively regulate thermal nociception, but the function of CCKLR-17D3 in nociceptive regulations resides elsewhere.

**Figure 4.**
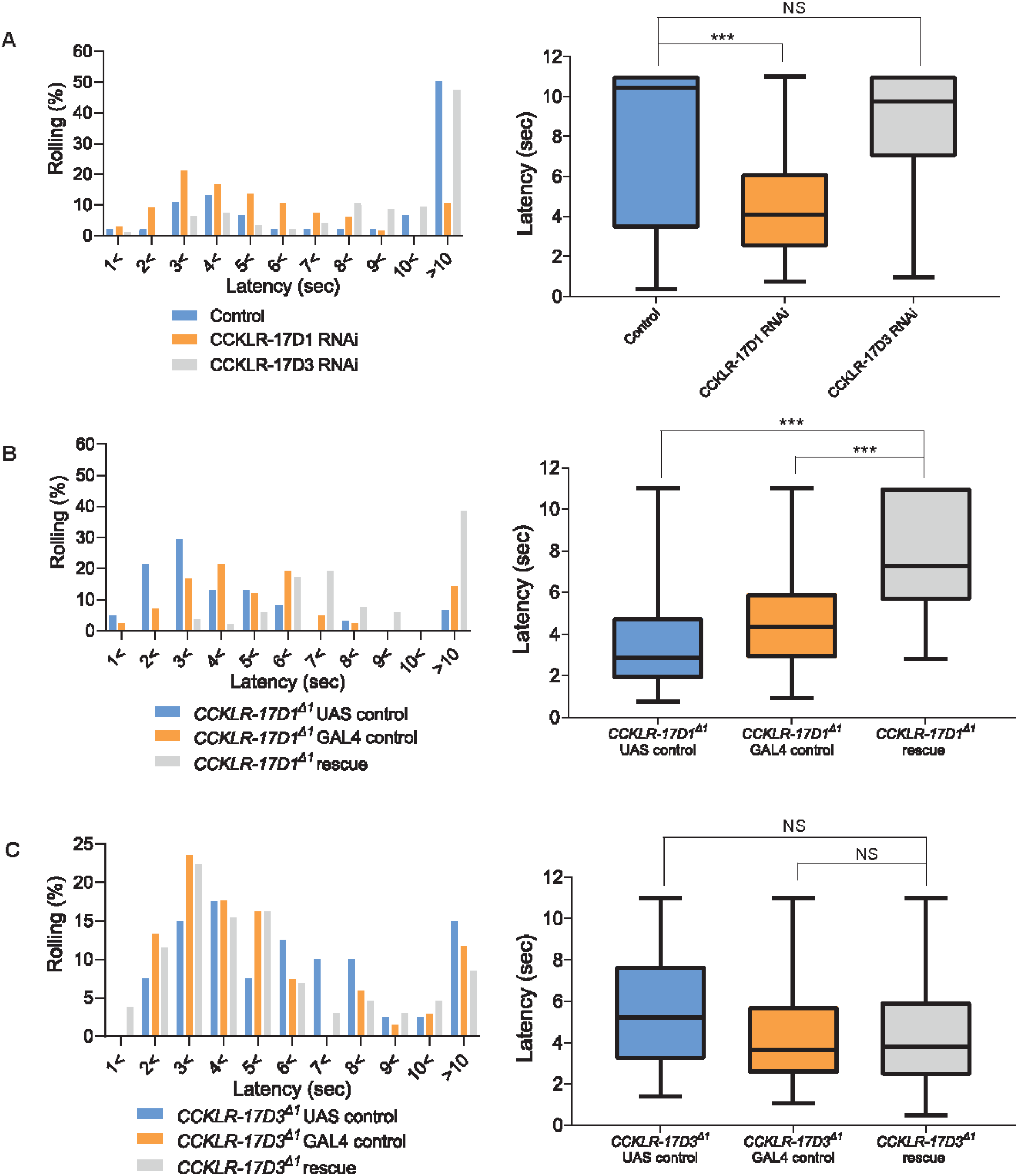
CCKLR-17D1 in Goro neurons is necessary and sufficient for normal thermal nociception. (A) RNAi of CCKLR-17D1 and CCKLR-17D3 using a Goro-GAL4 line *R69E06-GAL4*. Expressing CCKLR-17D1 RNAi (*R69E06-GAL4* x *yv; JF02644,* n = 66) but not CCKLR-17D3 RNAi (*R69E06-GAL4* x *yv; JF02968,* n = 95) with *R69E06-GAL4* caused significantly shorter latencies to 42 °C than controls (*R69E06-GAL4* x *yv; attp2,* n = 46). (Left) Histograms (Right) Box plots of latencies. *** p < 0.001, NS (non-significant) p > 0.05 Steel’s test. (B) Rescue of CCKLR-17D1 with *R69E06-GAL4*. The shortened latencies of *CCKLR-17D1^Δ1^* mutants observed in UAS controls (*CCKLR-17D1^Δ1^*; *UAS-CCKLR-17D1/+,* n = 61) or GAL4 controls (*CCKLR-17D1^Δ1^*; *R69E06-GAL4/+,* n = 42) were restored in the rescue genotype (*CCKLR-17D1^Δ1^*; *UAS-CCKLR-17D1/+; R69E06-GAL4/+,* n = 52). (Left) Histograms (Right) Box plots of latencies. *** p < 0.001 Steel’s test. (C) Rescue of CCKLR-17D3 with *R69E06-GAL4*. The shortened latencies of *CCKLR-17D3^Δ1^*mutants were unaltered in the rescue genotype (*CCKLR-17D3^Δ1^*; *UAS-CCKLR-17D3/+; R69E06-GAL4/+,* n = 130) compared with UAS controls (*CCKLR-17D3^Δ1^*; *UAS-CCKLR-17D3/+,* n = 40) or GAL4 controls (*CCKLR-17D3^Δ1^*; *R69E06-GAL4/+,* n = 68). (Left) Histograms (Right) Box plots of latencies. NS (non-significant) p > 0.05 Steel’s test. All box plots show median (middle line) and 25th to 75th percentiles with whiskers indicating the smallest to the largest data points.

Activation of Goro neurons elicits nocifensive rolling in larvae (53). Hence, if CCKLR-17D1 functions in Goro neurons to negatively regulate nociception, the lack of CCKLR-17D1 should sensitize these neurons to noxious heat. We directly addressed this hypothesis by using a Ca^2+^ imaging technique we had previously developed (32, 35), whereby we monitored GCaMP6m signals in Goro neurons while locally applying thermal ramp stimuli to the larval body wall. In our thermal ramp stimulation protocol, the temperature around the larval CNS hardly reached 28 °C during the heat ramp stimulations (27.0 ± 0.2 °C at the peak, n = 12), suggesting that the larval nociceptive system in the CNS that functions as an internal sensor for fast temperature increase (54) was unlikely to be activated.

Goro neurons in *CCKLR-17D1^Δ1^* mutants showed a significantly steeper GCaMP6m signal increase from 40 to 50 °C in comparison with the wild-type controls (Figure 5A and B, Video 1 and 2). Since the baseline fluorescence levels of GCaMP6m were not significantly different in the wild-type and *CCKLR-17D1^Δ1^* mutants (Figure 5C), these data demonstrate that Goro neurons in *CCKLR-17D1^Δ1^* mutants are specifically sensitized to a noxious range of heat. Suppressing CCKLR-17D1 by RNAi in Goro neurons also induced significantly sensitized responses of Goro to noxious temperatures of 44–49 °C (Figure 5D-F, Video 3 and 4). In contrast, Goro neurons in *CCKLR-17D3^Δ1^*mutants exhibited GCaMP6m signals that were mildly elevated but largely parallel compared with that of the controls (Figure 5-figure supplement 1, Video 5 and 6), providing further evidence for the major functioning of CCKLR-17D3 in nociceptive regulation outside Goro neurons. Considered together, these data demonstrate that CCKLR-17D1 functions to negatively regulate the activity of Goro neurons, thereby attenuating behavioral nociceptive responses.

**Figure 5.**
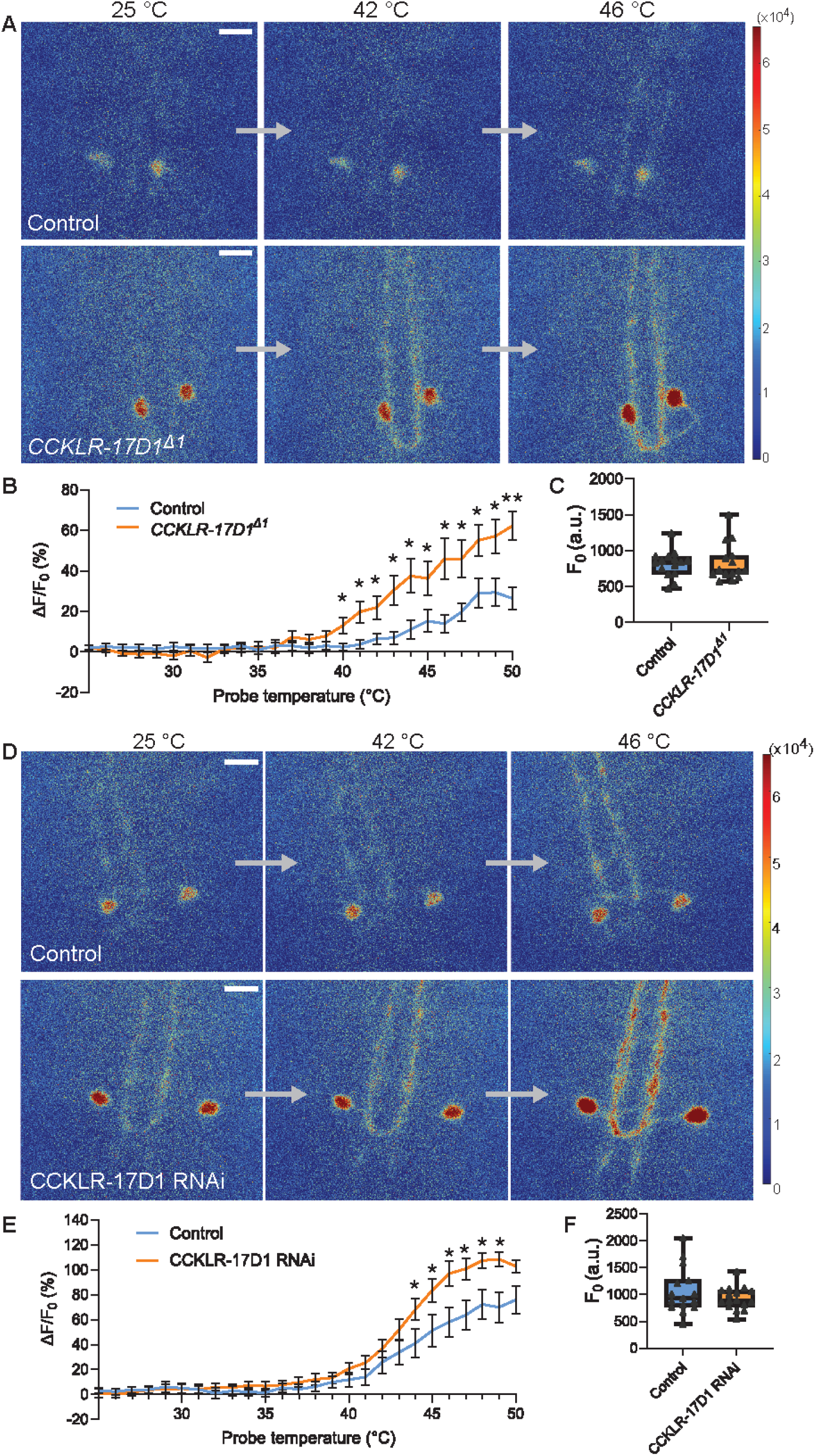
Goro neurons lacking CCKLR-17D1 show sensitized responses to noxious heat. (A) Representative still images showing thermal activation of Goro neurons in the controls (top, *yw*/Y; *R69E06-GAL4 UAS-GCaMP6m/+*) and *CCKLR-17D1^Δ1^* mutants (bottom, *CCKLR-17D1^Δ1^/Y; R69E06-GAL4 UAS-GCaMP6m*/+). See also Video 1 and 2. (B) Average percent increase of GCaMP6m fluorescence intensity relative to the baseline (ΔF/F_0_) during heat ramp stimulations in controls (n = 15) and *CCKLR-17D1^Δ1^* (n = 15). * p < 0.05, ** p < 0.01 Mann-Whitney’s U-test. (C) Basal GCaMP6m signal levels (F_0_) in controls (n = 15) and *CCKLR-17D1^Δ1^* mutants (n = 15). p > 0.6 Mann-Whitney’s U-test. (D) Representative stills showing thermal activation of the controls (top, *R69E06-GAL4 UAS-GCaMP6m* x *yv; attp2*) and Goro neurons expressing CCKLR-17D1 RNAi (bottom, *R69E06-GAL4 UAS-GCaMP6m* x *yv; JF02644*). See also Video 3 and 4. (E) Average percent increase of GCaMP6m fluorescence intensity relative to the baseline (ΔF/F_0_) during heat ramp stimulations in controls (n = 15) and CCKLR-17D1 RNAi (n = 16). * p < 0.05 Mann-Whitney’s U-test. (F) Basal GCaMP6m levels (F_0_) in controls (n = 15) and CCKLR-17D1 RNAi (n = 16). p > 0.6 Mann–Whitney’s U-test. Error bars represent standard error. All box plots show median (middle line) and 25th to 75th percentiles with whiskers indicating the smallest to the largest data points. All scale bars represent 20 µm.

### Neuronal projections and thermal responsiveness of DSK-expressing neurons

If CCKLR-17D1 is involved in regulating the activity of Goro neurons, how can DSK be conveyed from the brain to the VNC? We noticed that some of the *DSK-GAL4* positive brain neurons sent descending neural processes to the VNC (Figure 6A), which were all anti-FLRFa positive (Figure 6B). Furthermore, the anti-FLRFa signals in the descending projections were completely absent in the *dsk* mutants (Figure 6C), suggesting that these descending projections likely originated from the DSK-expressing brain cells, namely MP1 and/or Sv. In analyzing *DSK-2A-GAL4*, we found that MP1 neurons in fact sent the descending projections to the VNC (Figure 6D). To further understand the projection patterns of MP1 and Sv neurons in detail, we performed single-cell labeling using an FLP-out technique and revealed that MP1 neurons sent descending projections contralaterally to the VNC (Figure 6E), while Sv neurons projected within the brain (Figure 6F). In comparison with the longitudinal processes to the VNC, MP1 neural processes in the brain possessed few anti-FLRFa positive puncta, which represent DSK in these neurons (Figure 6G). Furthermore, when the somatodendritic marker *UAS-Denmark* (55) and the synaptic vesicle marker *UAS-syt::eGFP* were expressed in MP1 neurons, Denmark preferentially localized in the neural processes within the brain while syt::eGFP strongly accumulated in the processes descending to the VNC (Figure 6H). Collectively, these data demonstrate that MP1 neurons project DSK-positive descending axons to the VNC from the brain.

**Figure 6.**
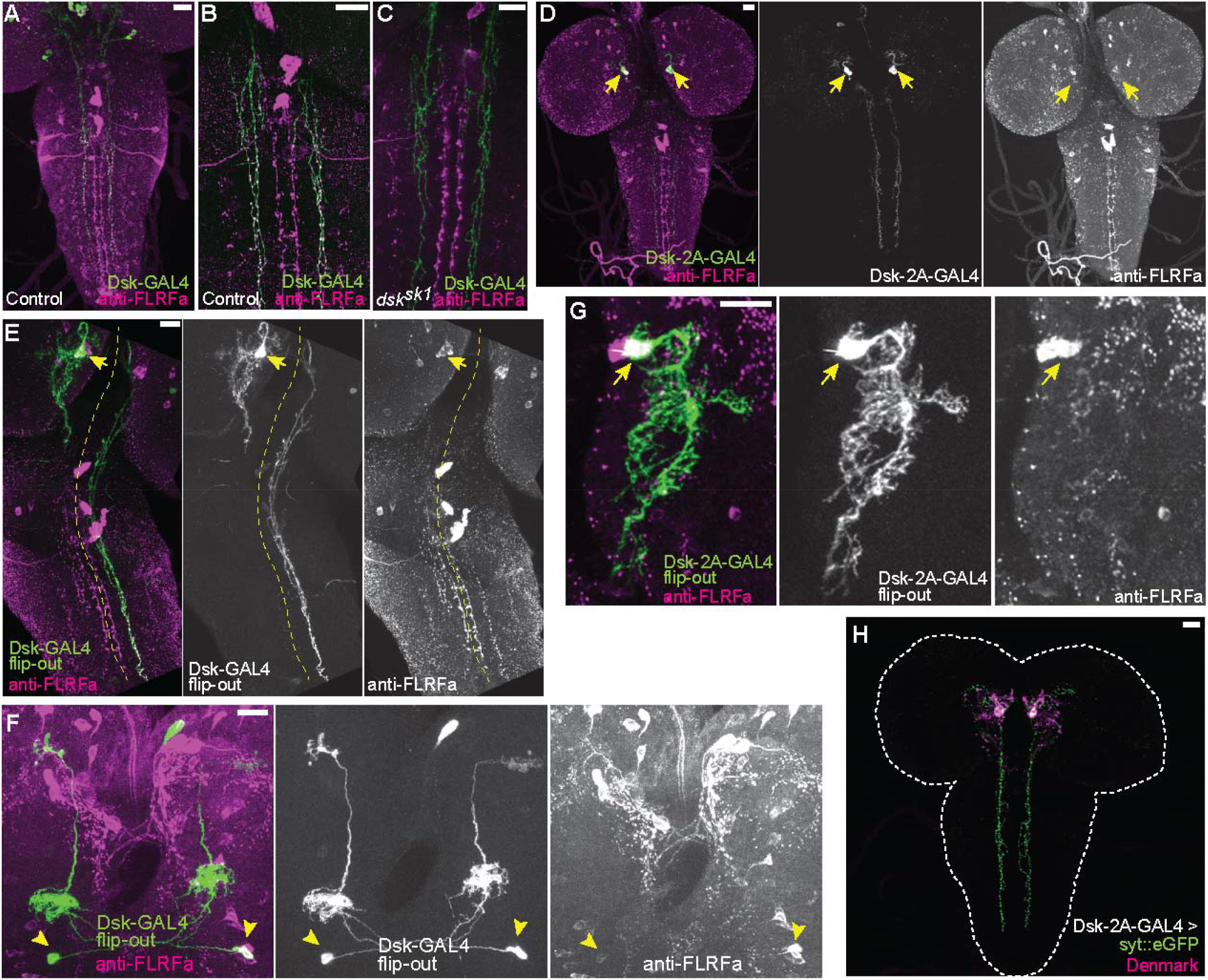
Projection patterns of DSK-expressing neurons in the larval CNS. (A) Representative image showing descending axons from *DSK-GAL4* positive brain neurons (green) to the larval VNC. (B) Representative image showing all descending projections labeled by *DSK-GAL4* (green) harbor punctate anti-FLRFa signals (magenta) in the wild-type. (C) Representative image showing the complete absence of punctate anti-FLRFa signals (magenta) in the descending projections labeled by *DSK-GAL4* (green) in the *dsk^sk1^* mutants. (D) Image showing the descending projections of MP1 neurons (arrows) marked by *DSK-2A-GAL4*. (E) Image showing a single FLP-out clone of MP1 neuron (arrow) contralaterally sending descending axons to the larval VNC. The yellow dashed line indicates the midline. (F) Image showing two FLP-out clones of Sv neurons (arrowheads) sending ascending axons to both sides of the brain. (G) A projection image showing the MP1 neurites in the brain. Note that few anti-FLRFa-positve puncta are associated with MP1 neurites within the brain. Similar expression patterns were observed in multiple samples (n = 3). The arrow indicates the anti-FLRFa-positve MP1 soma. (H) Representative image showing localizations of syt::eGFP (green) and Denmark (magenta) in MP1 neurons (*DSK-2A-GAL4* x *UAS-syt::eGFP UAS-Denmark*). Similar expression patterns were observed in multiple samples (n = 7). All scale bars represent 20 µm.

Next, we investigated whether some of these DSK-expressing neurons are responsive to noxious thermal stimuli. In order to examine the responsiveness of multiple neurons in intact larvae, we took the snapshot approach using CaMPARI, a genetically-encoded Ca^2+^ sensor whose fluorescence irreversibly changes from the green to the red region of the spectrum in response to irradiation with 405 nm UV light in a Ca^2+^ dependent manner, and is thus useful to monitor the activities of multiple neurons simultaneously in intact animals (56). When the larvae expressing CaMPARI2 (an improved version of CaMPARI (57)) in Goro neurons were stimulated by local thermal ramp stimuli under a 405 nm UV light, a significantly increased CaMPARI2 photoconversion was detected compared to the control group that had only been irradiated with 405nm UV (Figure 7A and B), confirming that by using CaMPARI2 in our experimental setup we managed to successfully capture the increased activity of Goro neurons in response to noxious heat in intact larvae. We then tested the responsiveness of *DSK-2A-GAL4* neurons using the same stimulation and photoconversion protocol and found that none of the MP1 or Sv neurons, nor IPCs showed increased CaMPARI2 photoconversion in response to local thermal ramp stimuli (Figure 7A, C-E). Interestingly, MP1 neurons exhibited high CaMPARI2 photoconversion regardless of the presence of local heat ramp stimuli, which was even comparable to that observed in Goro neurons activated by noxious heat stimuli (Figure 7A and C). These data suggest that none of the *DSK-2A-GAL4* neurons are clearly responsive to noxious heat, but the markedly high CaMPARI2 photoconversion observed in MP1 neurons raises the possibility that these neurons may be active regardless of the presence of noxious heat stimuli.

**Figure 7.**
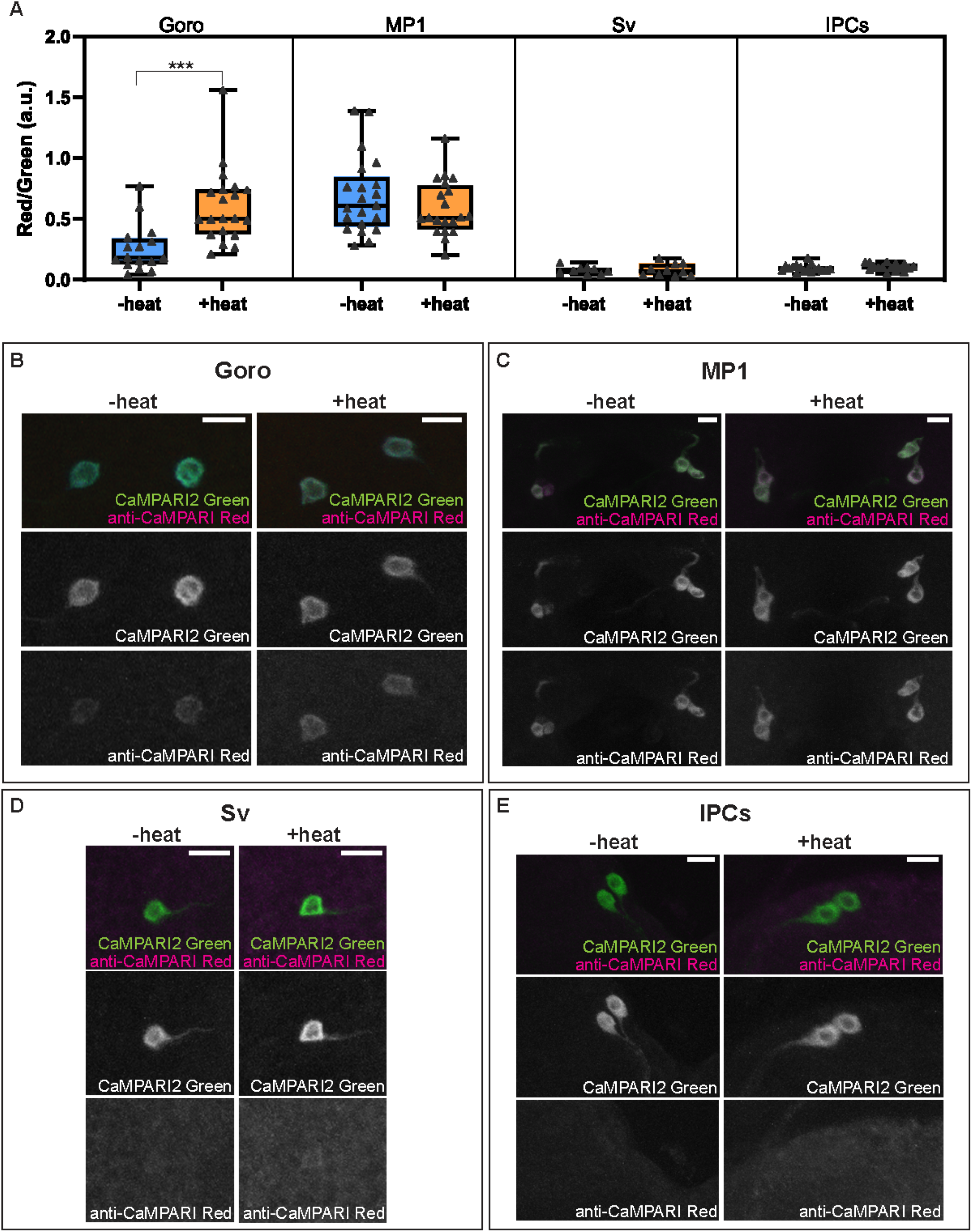
Activity and thermal responsiveness of DSK-expressing neurons. (A) Box plots of the quantified ratio of CaMPARI2 green signals and photoconverted CaMPARI2 red signals detected by anti-CaMPARI-red in Goro, MP1, Sv, and IPCs (Red/Green) with or without local heat ramp stimulation. Goro neurons showed a significantly higher Red/Green ratio of CaMPARI2 when local heat ramp stimulation was applied to the larvae (n = 16 and 20, p < 0.001, Mann-Whitney’s U-test). MP1 neurons showed a high Red/Green ratio of CaMPARI2 both with and without local heat stimulation (n = 21 and 20, p > 0.3, Mann-Whitney’s U-test). The Red/Green ratio of CaMPARI2 was low in Sv compared to MP1 neurons, and CaMPARI2 photoconversion was not increased by local heat stimulation (n = 9 and 10, p > 0.5, Mann-Whitney’s U-test). IPCs also exhibited low CaMPARI2 photoconversion regardless of the presence of local heat ramp stimulation (n = 17 and 16, p > 0.25, Mann-Whitney’s U-test). (B-E) Representative images showing CaMPARI2 green signals (green) and anti-CaMPARI-red signals (magenta) in Goro (B), MP1 (C), Sv (D), and IPCs (E). All box plots show median (middle line) and 25th to 75th percentiles, with whiskers indicating the smallest to the largest data points. Scale bars represent 10 µm.

### Connectivity between DSKergic and Goro neurons

To gain more insights into the connectivity between DSK-expressing brain neurons and Goro neurons, we further examined the anatomical relationship between MP1 axons and Goro neurons in the VNC. Performing double-labeling experiments, we found that MP1 axons with anti-FLRFa signals (representing DSK in MP1 axons) partially overlapped with Goro neurites in the thoracic segments of the larval VNC (Figure 8A). GFP reconstitution was reproducibly detected between MP1 and Goro neurons using the CD4-GRASP (GFP Reconstitution Across Synaptic Partners) system (58, 59), thus confirming that MP1 axons and Goro neurons are indeed in close proximity (Figure 8B). Since the CD4-GRASP is known to detect general cell-cell contacts (60, 61), we further investigated whether MP1 and Goro neurons are synaptically connected by using the nSyb-GRASP technique, which specifically enables synapse detection by localizing one of the split-GFP fragments in presynaptic sites (62). However, no signals of reconstituted GFP were detected between MP1 and Goro with nSyb-GRASP, although we found proximate localizations of nSyb-GFP_1-10_ in the MP1 axons with the Goro neurites expressing CD4::GFP_11_ (Figure 8C). We also employed the *trans*-Tango system (63) as another tool to detect synaptic connectivity. However, when the *trans*-Tango ligand was expressed by *DSK-2A-GAL4,* synaptic connectivity between MP1 and Goro was not indicated since the postsynaptic activation of *trans*-Tango signals was not observed in Goro neurons, although some *trans-*Tango-positive neurons were observed around Goro neurons (Figure 8D).

**Figure 8.**
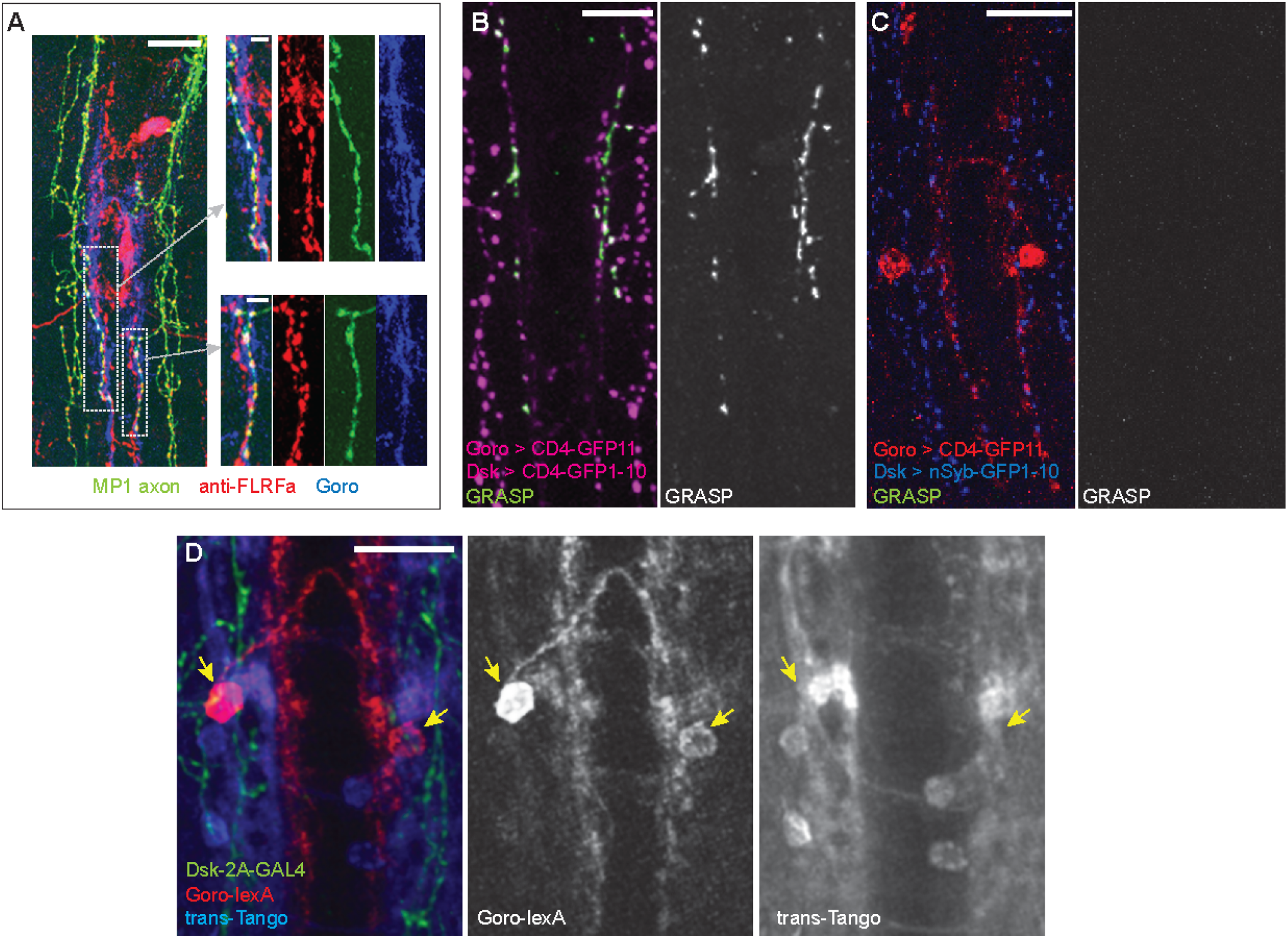
Synaptic connectivity between MP1 axons and Goro neurons is not detected. (A) Representative images of the larval VNC showing partially overlapping MP1 axons (green) and processes of Goro neurons (blue) that are associated with anti-FLRFa signals (red). MP1 axons partially overlapping Goro neurons were similarly observed in all examined samples (n = 5). (B) Representative image showing CD4-GRASP experiments between MP1 axons and Goro neurons (magenta). Signals of reconstituted GFP (GRASP, green) were detected in all examined samples (n = 5). (C) Representative image showing nSyb-GRASP experiments between MP1 axons (blue) and Goro neurons (red). Signals of reconstituted GFP (GRASP, green) were not detectable in any of the examined samples (n = 5). (D) Representative image showing *trans*-Tango experiments using *DSK-2A-GAL4* (*R69E06-lexA, lexAop-rCD2::RFP UAS-mCD8::GFP; DSK-2A-GAL4* x *UAS-myrGFP, QUAS-mtdTomato::3xHA; trans-Tango*). Transsynaptic activation of Tango signals (blue; mtdTomato::3xHA detected by anti-HA) was undetectable in Goro neurons (red; rCD2::RFP detected by anti-CD2). Green represents MP1 axons labeled by *DSK-2A-GAL4* (myrGFP and mCD8::GFP detected by anti-GFP). Signals of mtdTom::3xHA in Goro neurons were not detected in any of the examined samples (n = 4). Arrows indicate Goro somata. All scale bars represent 20 µm, except for the insets in (A) (5 µm).

Although we could not find positive evidence for synaptic connectivity between MP1 axons and Goro neurons, we further tested the hypothesis that DSK-expressing neurons in the brain may have functional connectivity to Goro neurons. To achieve this, we first attempted artificial activation experiments of *DSK-2A-GAL4* neurons using *UAS-dTRPA1* to determine whether activating DSK-expressing neurons actually attenuates larval nociception. dTRPA1 is a cation channel gated by warm temperature (> 29 °C) that has been used as a tool to artificially activate neurons of interest in *Drosophila* (64). When the larvae expressing dTRPA1 in C4da nociceptors with *ppk1.9-GAL4* were placed in a chamber at 35°C, dTRPA1-induced thermogenetic activation of C4da nociceptors caused nocifensive rolling responses within two seconds in the majority of animals, as reported previously (65) (Figure 9A and B). In contrast, when the larvae expressing dTRPA1 simultaneously in *ppk1.9-GAL4* and *DSK-2A-GAL4* were placed in a chamber at 35°C, the additional activation of *DSK-2A-GAL4* neurons to C4da nociceptors resulted in a markedly reduced percentage of larvae showing rolling responses within two seconds and in significantly lengthened latencies to exhibit a rolling response (Figure 9A and B). None of the animals expressing dTRPA1 with *DSK-2A-GAL4* alone showed a rolling response within 10 seconds (Figure 9A).

**Figure 9.**
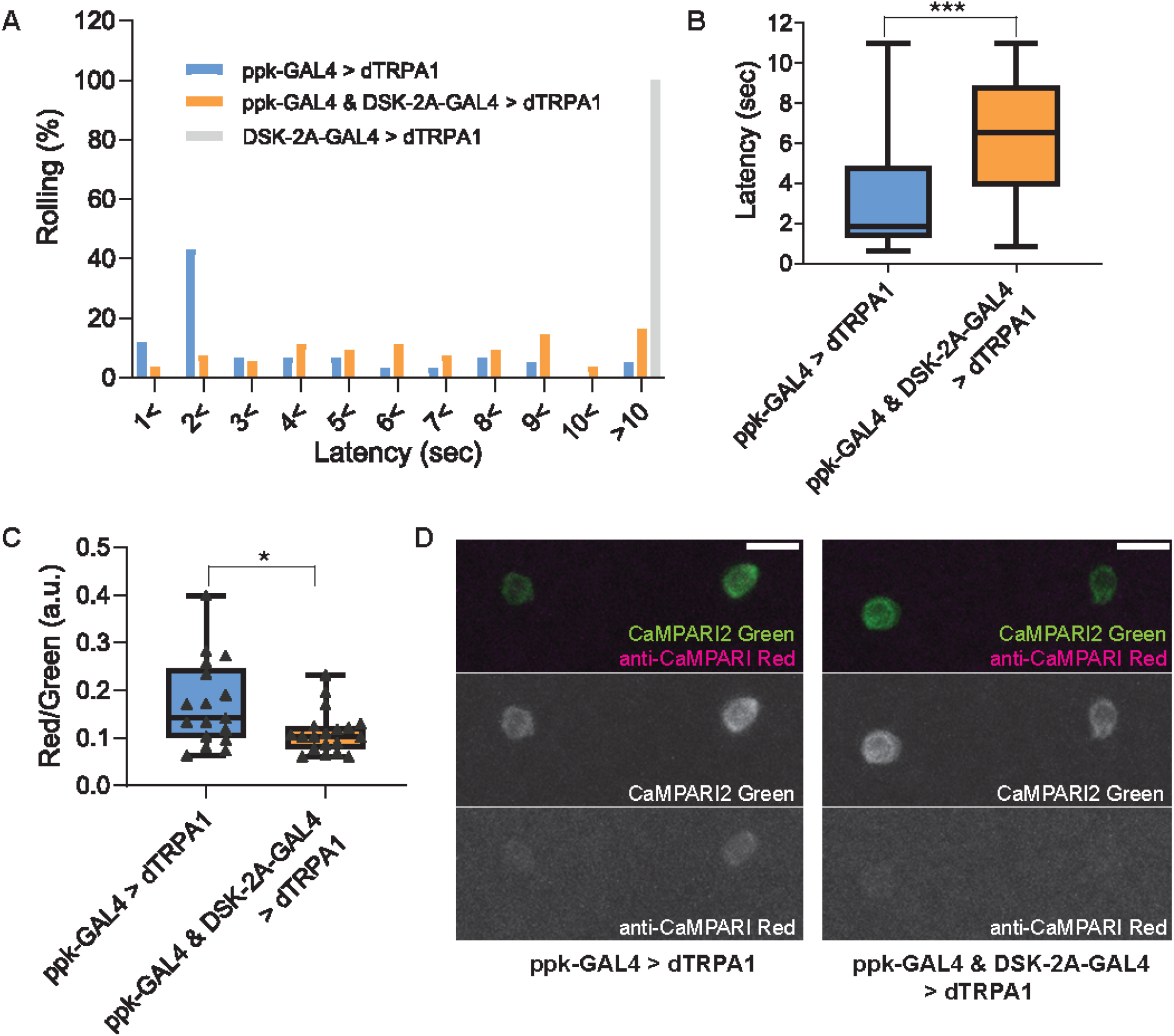
Thermogenetic activation of *DSK-2A-GAL4* neurons inhibits larval nociception and Goro neurons. (A and B) Thermogenetic activations of nociceptors and/or *DSK-2A-GAL4* neurons using *UAS-dTRPA1*. (A) Histogram representing the percentage of larvae exhibiting nociceptive rolling. Upon thermogenetic activations, 55% of larvae expressing dTRPA1 in nociceptors (ppk-GAL4 > dTRPA1; *ppk1.9-GAL4* x *UAS-dTRPA1*, n = 58) exhibited nociceptive rolling within 2 seconds. In contrast, only 11% of larvae with simultaneous activations of nociceptors and *DSK-2A-GAL4* showed rolling within 2 seconds (ppk-GAL4 & DSK-2A-GAL4 > dTRPA1; *ppk1.9-GAL4 DSK-2A-GAL4* x *UAS-dTRPA1*, n = 54). Thermogenetic activations of *DSK-2A-GAL4* neurons alone (DSK-2A-GAL4 > dTRPA1; *DSK-2A-GAL4* x *UAS-dTRPA1*, n = 40) did not trigger nociceptive responses even after 10 seconds. (B) Box plots showing latencies. Thermogenetic activation of *DSK-2A-GAL4* simultaneously with nociceptors caused a significantly longer latency to rolling responses (Mann-Whitney U-test, *** p < 0.001). (C and D) CaMPARI2 snapshot imaging of the activity of Goro neurons activated by thermogenetic stimulation of C4da nociceptors. (C) Box plots of the ratio of CaMPARI2 green signals and photoconverted CaMPARI2 red signals detected by anti-CaMPARI-red in Goro neurons (Red/Green). Compared to the larvae with thermogenetic activation of C4da nociceptors only (ppk-GAL4 > dTRPA1; *ppk1.9-GAL4, UAS-dTRPA1* x *R69E06-lexA; lexAop-CaMPARI2*; n = 17), the larvae with simultaneous activation of C4da and DSK-2A-GAL4 neurons (ppk-GAL4 & DSK-2A-GAL4 > dTRPA1; *ppk1.9-GAL4, UAS-dTRPA1; DSK-2A-GAL4* x *R69E06-lexA; lexAop-CaMPARI2*; n = 19) showed significantly decreased CaMPARI2 photoconversion in Goro neurons (Mann-Whitney U-test, * p < 0.025). (D) Representative images showing CaMPARI2 green signals (green) and anti-CaMPARI-red signals (magenta) in Goro neurons of animals with thermogenetic C4da activation (ppk-GAL4 > dTRPA1) and with simultaneous thermogenetic activation of C4da and DSK-2A-GAL4 neurons (ppk-GAL4 & DSK-2A-GAL4 > dTRPA1). All box plots show median (middle line) and 25th to 75th percentiles with whiskers indicating the smallest to the largest data points. Scale bars represent 10 µm.

The above data suggest that activation of *DSK-2A-GAL4* neurons inhibits larval nociception. We then investigated whether activating *DSK-2A-GAL4* neurons could attenuate the activity of Goro neurons, by combining the artificial activation experiments described above with CaMPARI2 snapshot imaging of Goro neurons. Indeed, CaMPARI2 photoconversion in Goro neurons induced by the thermogenetic activations of C4da nociceptors was significantly decreased by 35% with the simultaneous activation of *DSK-2A-GAL4* neurons (Figure 9C and D). Given that MP1 neurons were the only cells that were labeled by *DSK-2A-GAL4* with 100% reproducibility, while the expression of *DSK-2A-GAL4* in IPCs and Sv neurons was minor and stochastic (Figure 2F), the observed inhibition of behavioral nociception and Goro neurons caused by activating *DSK-2A-GAL4* neurons is mostly attributable to the activation of MP1 neurons. Thus, these data support the functional connectivity between DSKergic MP1 neurons in the brain and Goro neurons.

## Discussion

In this study, we have demonstrated that (1) DSK and its receptors CCKLR-17D1 and CCKLR-17D3 are involved in negatively regulating thermal nociception, (2) Two sets of brain neurons, MP1 and Sv, express DSK in the larval nervous system, (3) One of the DSK receptors (CCKLR-17D1) functions in Goro neurons in the VNC to negatively regulate thermal nociception, and (4) Thermogenetic activation of *DSK-2A-GAL4* neurons attenuates both larval nociception and the activity of Goro neurons. Based on these results, we propose that the DSKergic neurons regulating the activity of Goro neurons constitute a descending inhibitory pathway of nociception from the brain to the VNC in larval *Drosophila* (Figure 10). To our knowledge, our findings represent the first evidence of a descending mechanism modulating nociception from the brain in a non-mammalian species.

**Figure 10.**
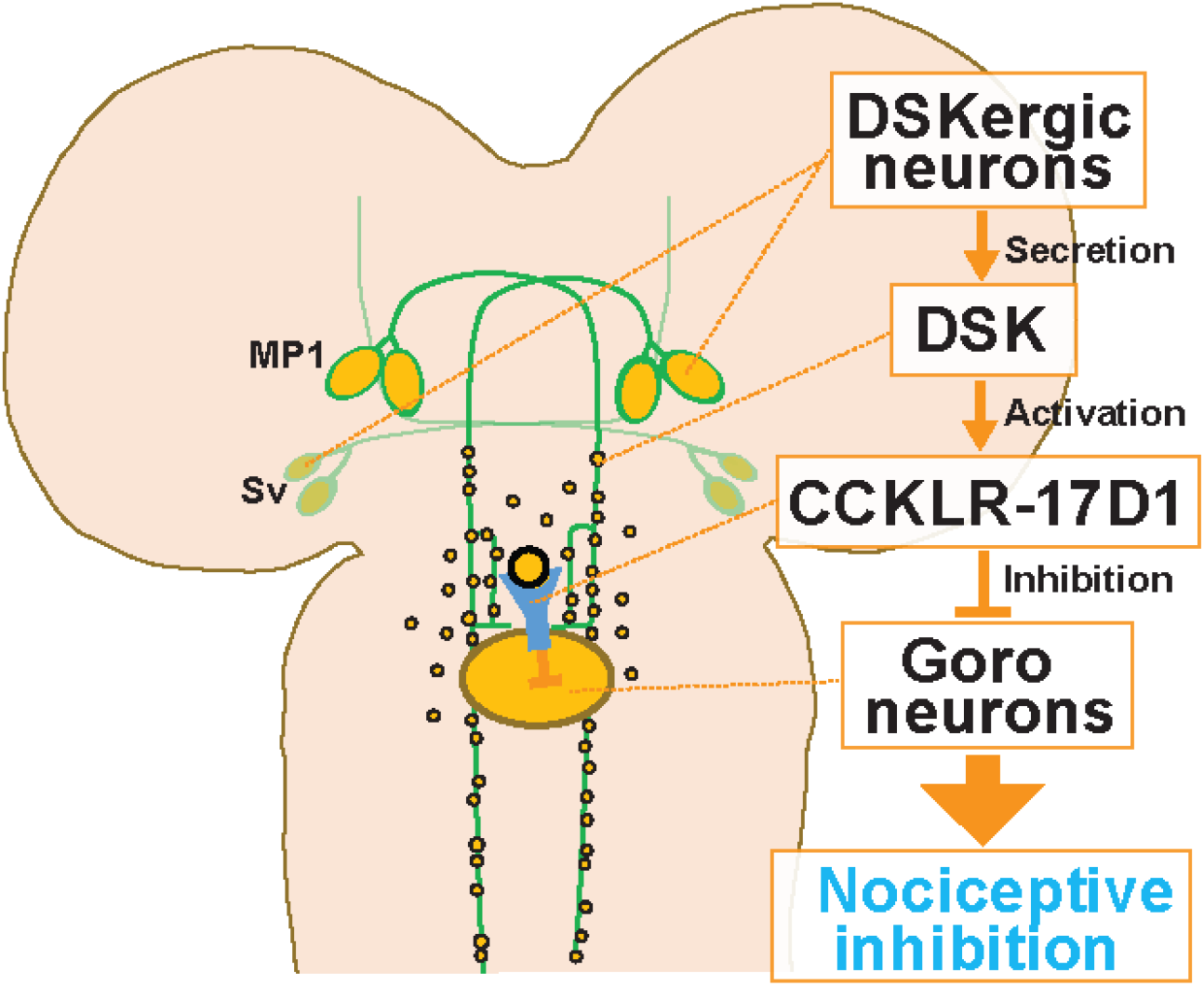
A schematic model of the DSKergic descending inhibitory pathway of nociception in larval *Drosophila*. DSK-expressing MP1 and/or Sv neurons in the brain secrete DSK peptides. DSK in the larval VNC activates the CCKLR-17D1 receptor expressed in Goro neurons, which subsequently inhibits the activity of Goro neurons, and ultimately larval nociceptive rolling responses.

### DSKergic descending system as a physiological modulator of nociception

DSK has been implicated in multiple physiological and developmental processes in *Drosophila* (19, 21–23). Previous studies have shown that *dsk* and *CCKLR-17D1* mutants exhibit significant reductions of synaptic growth and excitability in larval neuromuscular junctions (NMJ) and larval locomotion under bright light (45, 66), suggesting the importance of the DSK/CCKLR-17D1 signaling pathway in the developmental processes of motoneurons. However, in this study, no major developmental defects were observed in Goro neurons with CCKLR-17D1 RNAi (Figure 4-figure supplement 1E). Furthermore, the simultaneous thermogenetic activations of *DSK-2A-GAL4* neurons and C4da nociceptors inhibited larval nociception and the activity of Goro neurons (Figure 9). Given the pronociceptive role of Goro neurons, their reduced synaptic or neuronal activity should cause nociceptive insensitivity or reduced Ca^2+^ responses, which is contradicting to our observation that CCKLR-17D1 knockdown in Goro produced the thermal hypersensitivity and exaggerated Ca^2+^ responses (Figure 4 and 5). Thus, these data consistently support a physiological role for DSKergic descending signals in the modulation of the activity of Goro neurons rather than a developmental role, highlighting the functional differences of the DSK/CCKLR-17D1 pathway between the NMJ and the nociceptive system in the CNS.

### DSK in larval IPCs dispensable for nociceptive modulation

A previous study has reported the expression of DSK in a population of larval IPCs and its functions in responding to starving conditions (49). However, we observed that anti-FLRFa signals in IPCs persisted in *dsk* null mutants and that anti-FLRFa signals in IPCs hardly overlap with either *DSK-GAL4* or *DSK-2A-GAL4* expressions (Figure 2 and Figure 2-figure supplement 1). Although we used a different antibody from that in the study mentioned above, the number of IPCs visualized by anti-FLRFa was comparable to that of cells visualized by anti-DSK (Figure 2-figure supplement 1) (49). Regarding the functions of IPCs in nociception, Im et al. showed that silencing *dilp2-GAL4* positive IPCs has no significant effect on the baseline nociceptive responses (67). In our CaMPARI2 imaging, IPCs marked by *DSK-2A-GAL4* showed very low baseline activity and no responsiveness to noxious heat (Figure 7). Thus, the current and previous studies strongly suggest that IPCs are likely irrelevant for DSK-mediated nociceptive regulation, at least under normal conditions, although it is still possible for DSK to be expressed stochastically in a limited population of IPCs.

### Functional mechanisms of DSKergic nociceptive inhibitory system

Although our data suggest the existence of a DSKergic descending inhibitory system from the brain to the Goro neurons in the VNC, there are multiple remaining questions for future studies on the mechanisms by which DSK-expressing neurons in the brain regulate Goro neurons: First, whether both MP1 and Sv neurons are involved in nociceptive inhibition remains unclear. Our findings consistently point to MP1 neurons over Sv neurons as a major source of DSK regulating Goro neurons. However, given the potential of DSK for distant actions through diffusion, the possibility of Sv neurons communicating with Goro neurons cannot be eliminated. Further analysis using finer genetic/neuronal tools will be necessary to tease the functions of MP1 and Sv neurons apart and understand the functional mechanism of the DSKergic system in the regulation of larval nociception.

Second, how this DSKergic system is activated to regulate nociception *in vivo* needs further investigation. We observed very high CaMPARI2 photoconversion in MP1 neurons without noxious heat stimuli, which did not further increase in response to noxious heat application (Figure 7). These data raise the hypothesis that MP1 neurons may function as a tonically active brake for nociceptive circuits, rather than a simple negative feedback system that is activated by nociceptive stimuli as input. The tonically active model of MP1 function is also consistent with our data showing that *dsk* and *CCLR-17D1* mutations, as well as CCLR-17D1 RNAi in Goro neurons, all caused thermal hypersensitivity under normal conditions (Figure 1 and 4). Up to this point, no external or internal signals have been directly shown to be the input into larval DSKergic neurons. However, since DSK has been implicated in the regulation of several physiological processes such as hunger and stress, sensory or molecular signals that are involved in these physiological processes have the potential to serve as upstream cues that activate DSKergic neurons. Alternatively, it is also possible that MP1 neurons may have spontaneous activity. Further physiological studies on the activity and responsiveness of DSKergic neurons would be required to elucidate how the DSKergic system functions to regulate nociception in larvae under normal conditions.

Third, the transmission mechanisms of DSK from the brain to Goro neurons need further clarification. Our nSyb-GRASP and *trans-*Tango experiments consistently showed negative results, indicating no synaptic connectivity between MP1 descending axons and Goro neurons (Figure 8). Thus, the interaction between DSK from DSKergic neurons in the brain and Goro neurons in the VNC may be mediated non-synaptically through volume transmission, as described in many neuropeptidergic systems (68). Many of the CCKLR-expressing neurons in the VNC are distantly located from the descending MP1 axons (Figure 3 and 6). Since it is unlikely that all these CCKLR-expressing neurons are synaptically connected to MP1 axons, neuronal communications through volume transmission can be fairly assumed for DSKergic systems in the larval VNC. More detailed analyses at an electron microscopic level of the circuitry connectivity between MP1 and Goro neurons and of the localization of DSK-containing vesicles in DSKergic neurons as well as DSK receptors in Goro neurons would be necessary to further clarify the transmission mechanisms of DSK from the brain to Goro neurons.

### Multiple molecular pathways for DSK signaling in the regulation of nociception

The data presented in this study also suggest that DSK signaling could regulate larval nociception through multiple pathways other than the DSK/CCKLR-17D1 system. For example, while *CCKLR-17D3* mutants exhibited as severe thermal hypersensitivity as *CCKLR-17D1* mutants (Figure 1D and F), our RNAi and rescue experiments failed to locate the function of CCKLR-17D3 to Goro neurons (Figure 4A and C). Our GCaMP Ca^2+^ imaging also revealed that Goro neurons lacking CCKLR-17D3 showed modestly sensitized responses to noxious heat (Figure 5-figure supplement 1). These results suggest the major functioning of CCKLR-17D3 in Goro-independent nociceptive pathways. We also observed that DSK receptor mutants exhibited more severe thermal phenotypes than *dsk* mutants (Figure 1), which may indicate the presence of unidentified DSK- or CCKLR-dependent signaling pathways in nociception-related cells. Further research is apparently necessary to reveal the whole picture of nociceptive regulations mediated by DSK and its receptors in larval *Drosophila*.

### Potential conservation of CCK-mediated descending nociceptive controls

The CCK system is thought to be one of the most ancient neuropeptide systems, suggested to have multiple common physiological functions across taxa (16–20). In this study, we demonstrate that CCKergic signaling in *Drosophila* participates in nociceptive modulation through a descending inhibitory pathway similarly to the mammalian CCK system, adding new evidence about the conserved physiological roles of CCK.

Unlike the peripheral nociceptive systems, it is still challenging to align the *Drosophila* and mammalian CCKergic descending pathways due to low homologies of the CNS structures between *Drosophila* and mammals, wide-spread CCK expression in the mammalian CNS, and the multiple roles of the CCKergic systems in mammalian nociceptive controls (10–14, 69–73). However, the common usage of an orthologous molecular pathway in descending controls of nociception between the two evolutionarily distant clades raises a fascinating new hypothesis that the descending control from the brain may also be an ancient, conserved mechanism of nociception, which has emerged in the common ancestor of protostomes and deuterostomes. It will be of interest for future research to investigate whether the role of CCK signaling in descending nociceptive controls is also present in other species.

### Potentials of non-mammalian models to study the mechanisms of pain modulation and pain pathology in the CNS

Non-mammalian model systems have been increasingly recognized as powerful tools to identify novel pain-related molecular pathways (2, 74–77). However, the utilization of these models has so far been mostly limited to the research on peripheral pain pathophysiology, and few studies have used them to investigate central pain pathophysiology (2, 78). Descending nociceptive control mechanisms are crucial for central pain modulation and have been implicated in the development of chronic pain states in humans (5, 6). Thus, the current study opens the door to a new approach to using powerful neurogenetic tools and the simpler nervous system of *Drosophila* for elucidating the functional principles of descending nociceptive systems, which may potentially contribute to our understanding of the mechanisms underlying central pain modulation and pain pathology due to dysfunctions of descending modulatory pathways.

## Acknowledgments

We thank the NIG-Fly Stock Center, TRiP at Harvard Medical School (NIH/NIGMS R01-GM084947), the Bloomington Stock Center, and Drs. Masayuki Koganezawa, Takeshi Awasaki, Barry Ganetzky, and Marta Zlatic for fly stocks. We are grateful to Dr. Eve Marder for kindly providing the anti-FLRFa antibody. We also thank Drs. Yoshiki Hayashi, Makoto Hayashi, and Satoru Kobayashi for their support in imaging with Leica SP5 and SP8. This study was supported by grants from JSPS KAKENHI to K Honjo (17K14928, 19K06935, and 22K06328) and HT (17H01378 and 26250001), and grants from the National Center for Geriatrics and Gerontology (21–47), the Uehara Memorial Foundation, and the Mochida Memorial Foundation for Medical and Pharmaceutical Research to K Honjo.

## Author contributions

K Honjo conceived the research. IO, K Hashimoto, AY, MH, and K Honjo performed experiments and analyzed data. SK and HT generated dsksk1 mutant and DSK receptor GAL4 lines. K Honjo wrote the manuscript with input and edits from IO, SK, HT, and KFT.

## Supplemental figures

**Figure 1-figure supplement 1.**
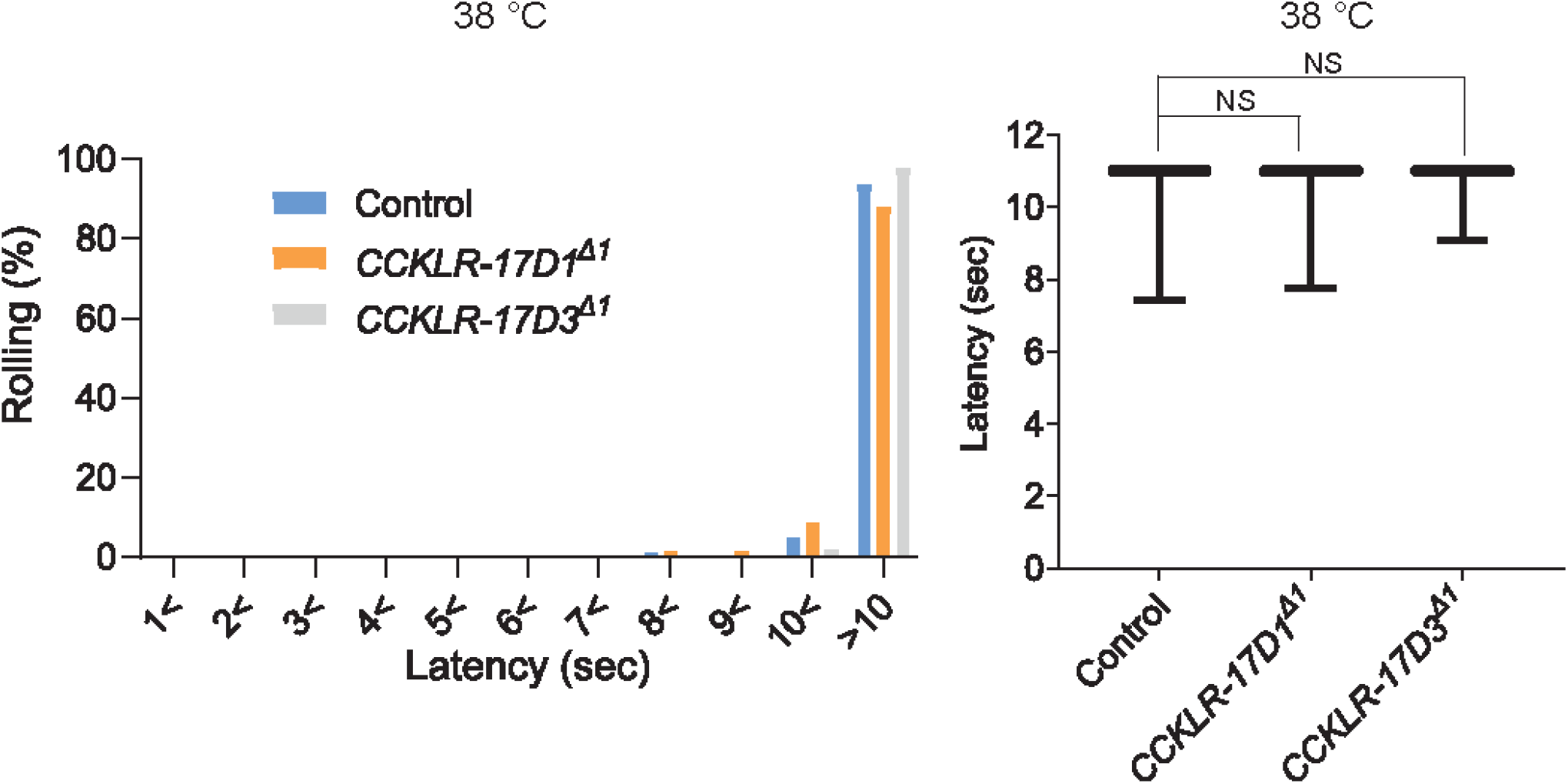
Thermal nociceptive thresholds in DSK receptor mutants are largely normal. Thermal responses of *CCKLR-17D1* and *CCKLR-17D3* mutants to a 38 °C probe. Both *CCKLR-17D1^Δ1^* (n = 58) and *CCKLR-17D3^Δ1^* (n = 48) showed comparable distributions of responding latencies to the controls (*yw*, n = 79). Box plots show median (middle line) and 25th to 75th percentiles with whiskers indicating the smallest to the largest data points. p > 0.4 Steel’s test.

**Figure 2-figure supplement 1.**
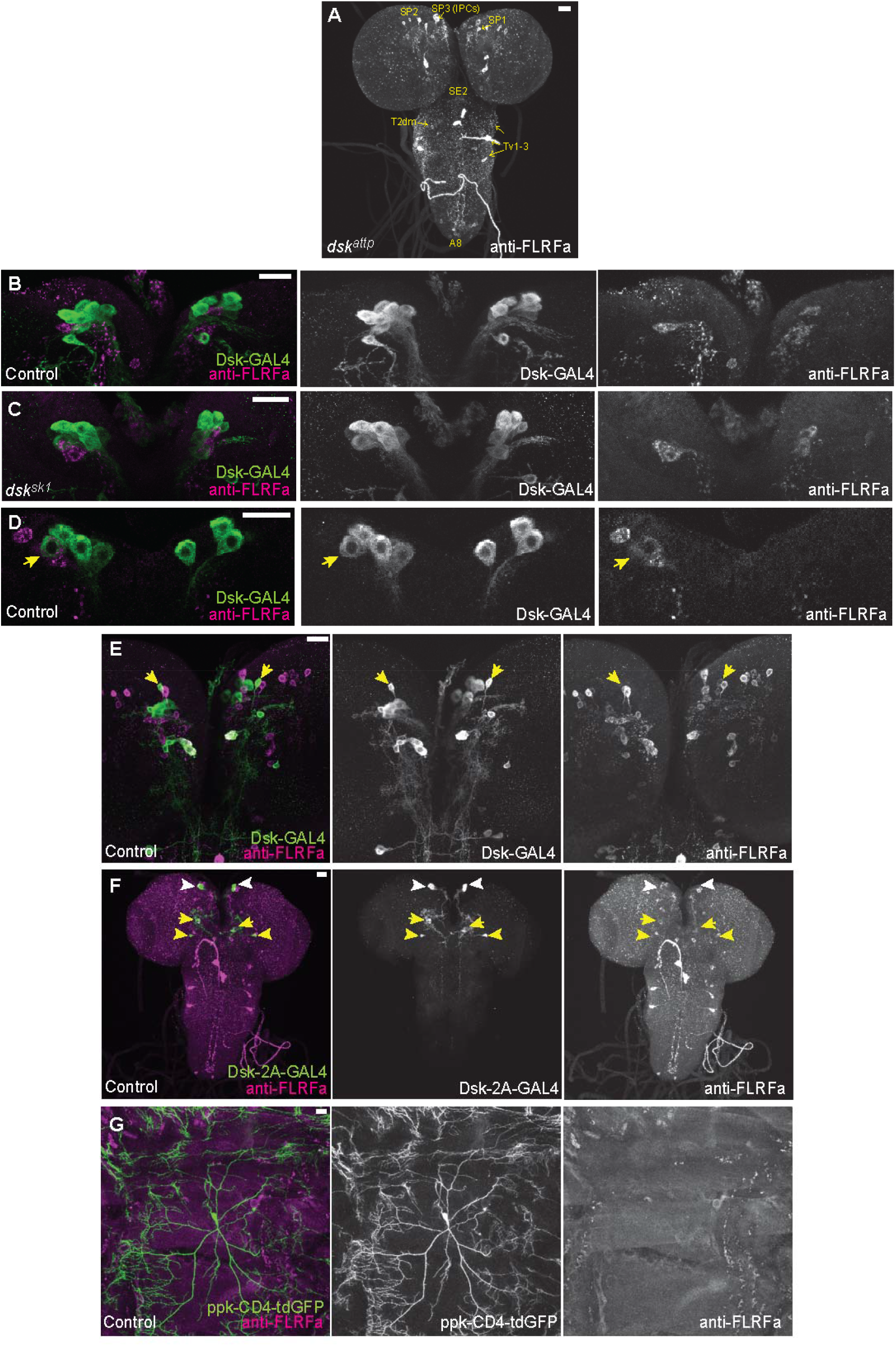
DSK expressions in the other brain neurons and peripheral sensory neurons. (A) Representative image showing staining of anti-FLRFa in *dsk^attp^*mutants. The staining patterns were indistinguishable from *dsk^sk1^*mutants. (B) Representative image showing staining patterns of *DSK-GAL4* and anti-FLRFa in the wild-type larval IPCs. In the wild-type brain, typically 4 to 7 IPCs are visualized by *DSK-GAL4* and 1 to 3 by anti-FLRFa consistently to the previous study by Soderberg et al. [49] that used anti-DSK raised by Nichols et al. [47] In contrast, few IPCs co-labelled were found, which is also somewhat consistent with the variability of DSK immunostaining in IPCs reported by the previous study [49]. (C) Representative image showing staining patterns of *DSK-GAL4* and anti-FLRFa in the *dsk^sk1^* larval IPCs. Comparable staining patterns of *DSK-GAL4* and anti-FLRFa were observed in the mutant IPCs. (D) An example of rare IPC that was co-labeled by *DSK-GAL4* and anti-FLRFa in the wild-type brain (arrow). (E) Representative image showing SP1 neurons (arrows) that were rarely co-labeled by *DSK-GAL4* and anti-FLRFa. Note that faint anti-FLRFa staining was only detected in the SP1 neuron in the right hemisphere. (F) An example image showing *DSK-2A-GAL4* expression in IPCs (white arrowheads), MP1 (yellow arrows), and Sv neurons (yellow arrowheads). (G) Representative image showing double-labeling of Class IV md neurons (*ppk-CD4::tdGFP*) and anti-FLRFa staining in the dorsal larval body wall. All scale bars represent 20 µm.

**Figure 3-figure supplement 1.**
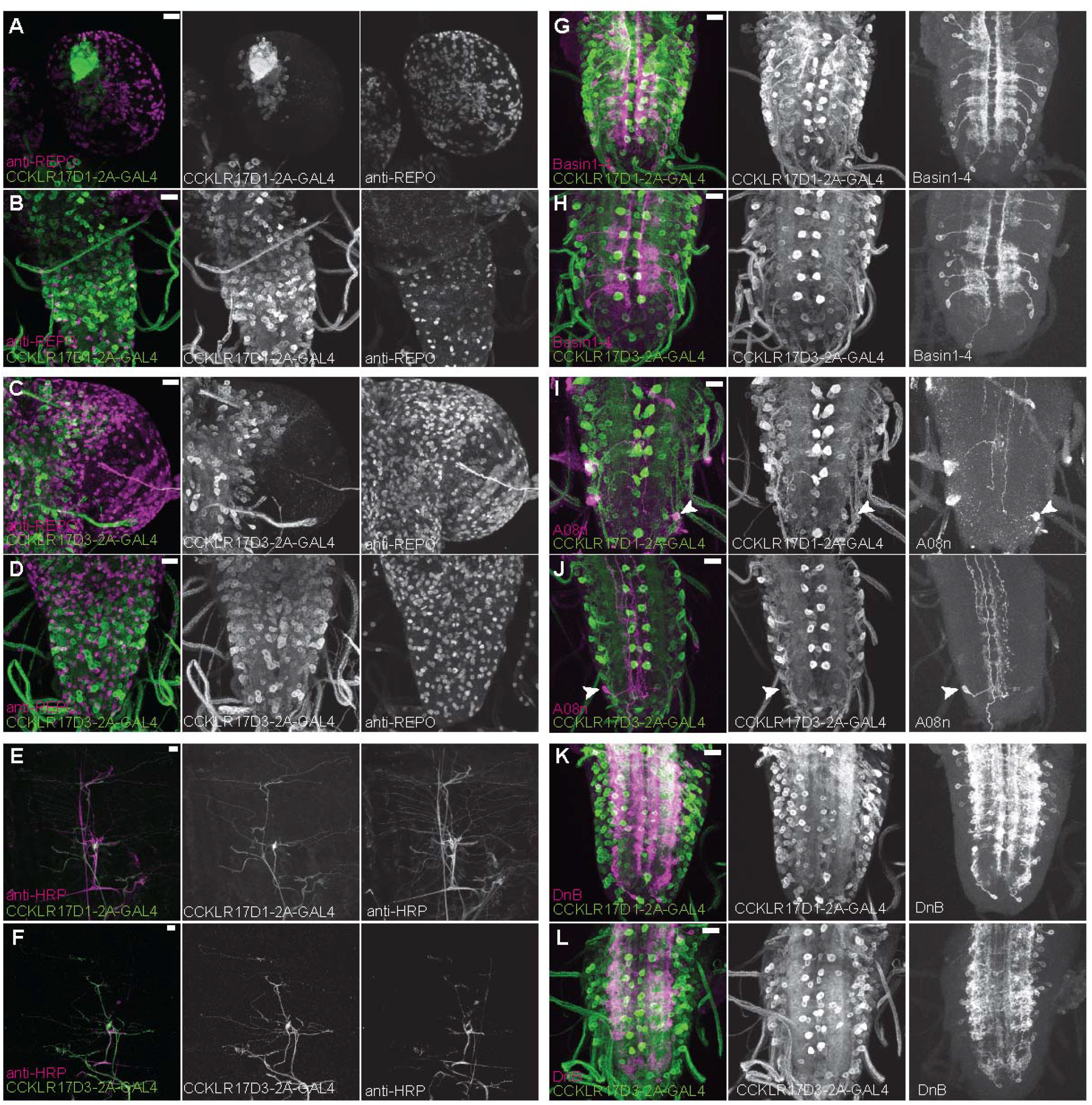
Expression patterns of DSK receptors in larval glial cells, peripheral tissue, and nociceptive interneurons. (A and B) Representative images showing double-labeling of *CCKLR-17D1-T2A-GAL4* and a glial cell marker anti-REPO in the larval brain (A) and the larval VNC (B), representing little expressions of *CCKLR-17D1-T2A-GAL4* in glial cells. (C and D) Representative images showing double-labeling of *CCKLR-17D3-T2A-GAL4* and a glial cell marker anti-REPO in the larval brain (C) and in the larval VNC (D), representing little expressions of *CCKLR-17D3-T2A-GAL4* in glial cells. (E and F) Representative images showing the expressions of *CCKLR-17D1-T2A-GAL4* (E) and *CCKLR-17D3-T2A-GAL4* (F) with a peripheral neuronal marker anti-HRP in the larval body wall, showing the absence of both GAL4s from peripheral sensory neurons other than es cells. (G and H) Expressions of *CCKLR-17D1-T2A-GAL4* (G) and *CCKLR-17D3-T2A-GAL4* (H) were negligible in Basin1 to 4 neurons labeled by *R72F11-lexA* in the larval VNC. (I and J) Negligible expression of *CCKLR-17D1-T2A-GAL4* (I) and *CCKLR-17D3-T2A-GAL4* (J) found in A08n neurons labeled by *R82E12-lexA* (arrowheads) in the larval VNC. (K and L) Expressions of *CCKLR-17D1-T2A-GAL4* (K) and *CCKLR-17D3-T2A-GAL4* (L) were negligible in DnB neurons labeled by *R70F01-lexA* in the larval VNC. All scale bars represent 20 µm.

**Figure 4-figure supplement 1.**
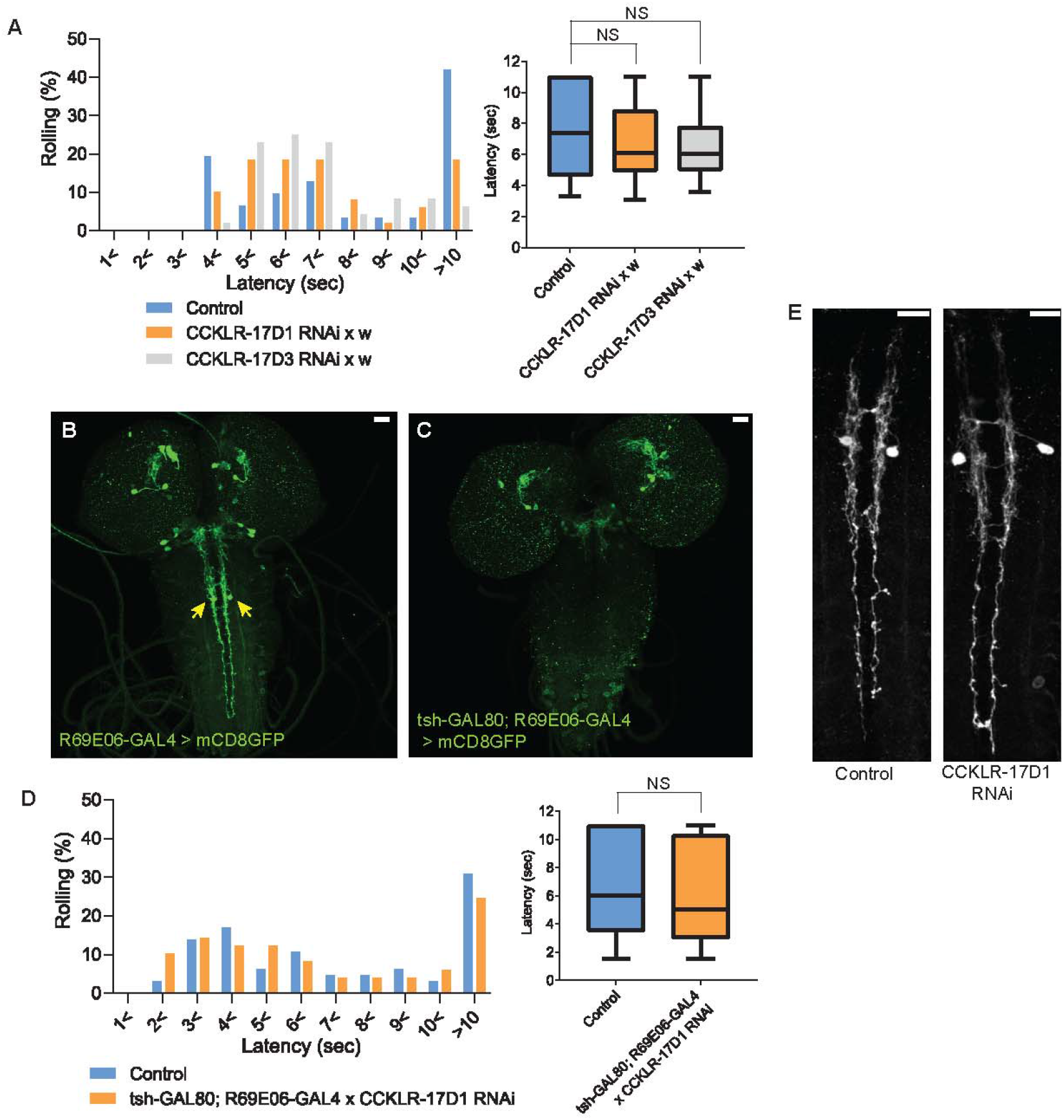
CCKLR-17D1 RNAi in Goro neurons induces thermal hypersensitivity in a GAL4-dependent manner, without affecting the morphology. (A) Without the GAL4 driver, UAS-CCKLR-17D1 RNAi (*yv;JF02644* x *w^1118^*, n = 49) and UAS-CCKLR-17D3 RNAi (*yv;JF02968* x *w^1118^*, n = 48) both did not cause thermal hypersensitivity to 42 °C compared with the control (*R69E06-GAL4* x *yv; attp2,* n = 31). p > 0.38 Steel’s test. (B and C) *tsh-GAL80* eliminates the expression of *R69E06-GAL4* from Goro neurons. The expression of *R69E06-GAL4* in Goro neurons (B) (*10xUAS-IVS-mCD8GFP; R69E06-GAL4* x *yv; attp2*) was eliminated when *tsh-GAL80* was combined with *R69E06-GAL4* (C) (*tsh-GAL80; R69E06-GAL4* x *40xUAS-IVS-mCD8GFP*). (D) When the expression of UAS-CCKLR-17D1 RNAi was eliminated from Goro neurons (*tsh-GAL80; R69E06-GAL4* x *yv; JF02644*, n = 49), larval nociceptive responses to a 42 °C probe was indistinguishable from the control (*R69E06-GAL4* x *yv; attp2,* n = 65). p > 0.33 Mann-Whitney’s U-test. (E) Goro neurons expressing CCKLR-17D1 RNAi (*10xUAS-IVS-mCD8GFP; R69E06-GAL4* x *yv; JF02644*) did not alter their number, position, and gross projection patterns in comparison with the control (*10xUAS-IVS-mCD8GFP; R69E06-GAL4* x *yv; attp2*). All scale bars represent 20 µm. All box plots show median (middle line) and 25th to 75th percentiles with whiskers indicating the smallest to the largest data points.

**Figure 5-figure supplement 1.**
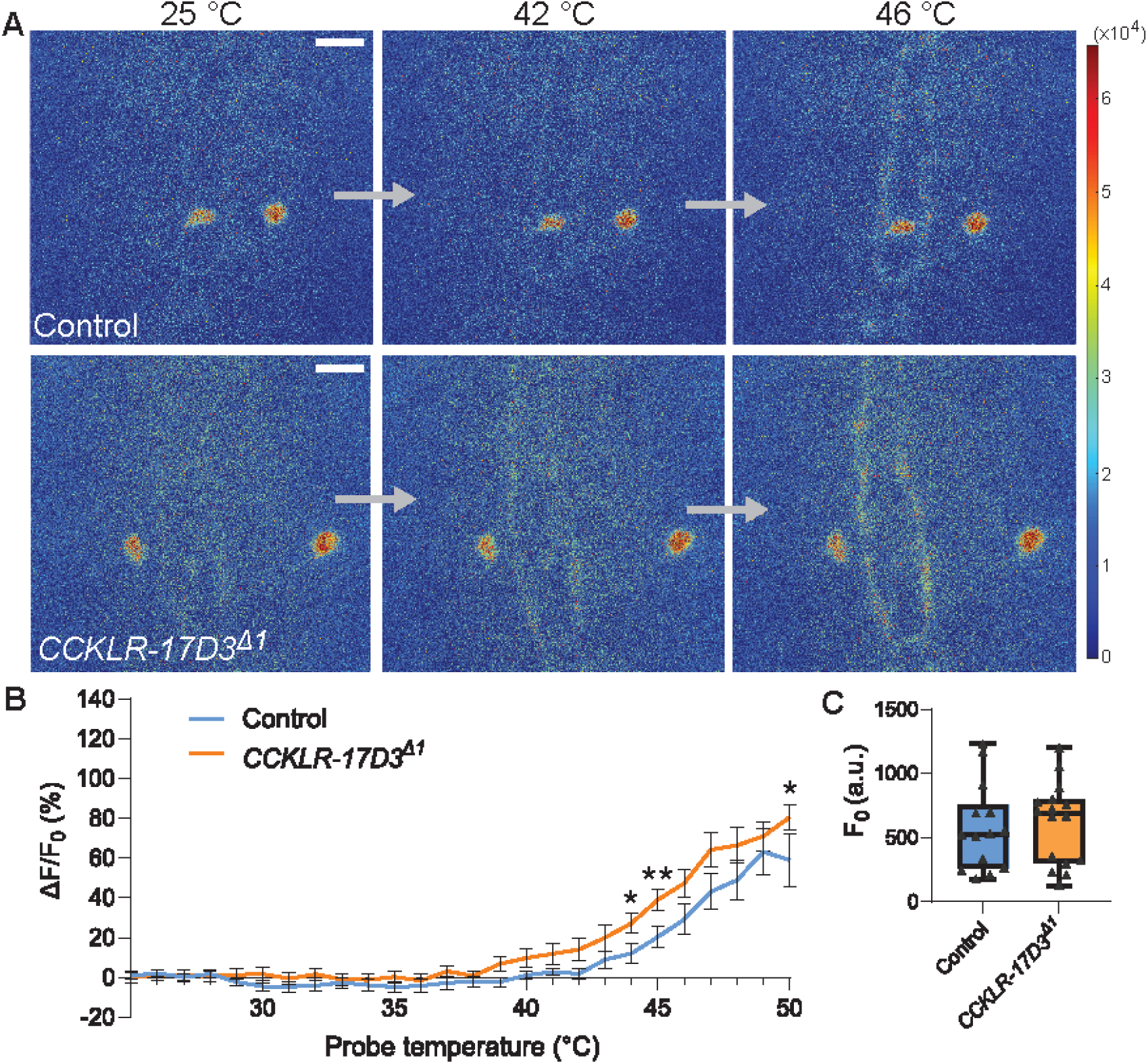
Thermal responsiveness of Goro neurons in CCKLR-17D3Δ1. (A) Representative still images showing thermal activation of Goro neurons in control animals (top, *yw*/Y; *R69E06-GAL4 UAS-GCaMP6m/+*) and *CCKLR-17D3^Δ1^* mutants (bottom, *CCKLR-17D3^Δ1^/Y; R69E06-GAL4 UAS-GCaMP6m*/+). See also Movie S5 and S6. Scale bars represent 20 µm. (B) Average percent increase of GCaMP6m fluorescence intensity relative to baseline (ΔF/F_0_) during heat ramp stimulations in *CCKLR-17D3^Δ1^* experiments. ΔF/F_0_ is plotted to binned probe temperature (interval = 1 °C). In comparison with controls, Goro neurons of *CCKLR-17D3^Δ1^* exhibited mildly elevated fluorescent increase of GCaMP6m, which reached statistical significance only at 44°C, 45°C, and 50°C. n = 14 and 16 for controls and *CCKLR-17D3^Δ1^*, respectively. * p < 0.05, ** p < 0.01 (Mann–Whitney’s U-test). Error bars represent standard errors. (C) Basal GCaMP6m signal levels (F_0_) did not differ between controls and *CCKLR-17D3^Δ1^*(n = 14 and 16). p > 0.5 (Mann–Whitney’s U-test). Box plots show median (middle line) and 25th to 75th percentiles with whiskers indicating the smallest to the largest data points.

**Video 1.**

Ca2+ imaging of Goro neurons in a control animal used for the CCKLR-17D1Δ1 experiments. See also Figure 5. The movie was generated from heat-mapped time-series projection images of GCaMP6m fluorescence using MATLAB. Time and probe temperature are indicated at the top left and top right corners, respectively.

**Video 2.**

Ca2+ imaging of Goro neurons in a CCKLR-17D1Δ1 mutant animal. See also Figure 5. The movie was generated from heat-mapped time-series projection images of GCaMP6m fluorescence using MATLAB. Time and probe temperature are indicated at the top left and top right corners, respectively.

**Video 3.**

Ca2+ imaging of Goro neurons in a control animal used for the CCKLR-17D1 RNAi experiments. See also Figure 5. The movie was generated from heat-mapped time-series projection images of GCaMP6m fluorescence using MATLAB. Time and probe temperature are indicated at the top left and top right corners, respectively.

**Video 4.**

Ca2+ imaging of Goro neurons in a CCKLR-17D1 RNAi animal. See also Figure 5. The movie was generated from heat-mapped time-series projection images of GCaMP6m fluorescence using MATLAB. Time and probe temperature are indicated at the top left and top right corners, respectively.

**Video 5.**

Ca2+ imaging of Goro neurons in a control animal used for the CCKLR-17D3Δ1 experiments. See also Figure 5-figure supplement 1. The movie was generated from heat-mapped time-series projection images of GCaMP6m fluorescence using MATLAB. Time and probe temperature are indicated at the top left and top right corners, respectively.

**Video 6.**

Ca2+ imaging of Goro neurons in a CCKLR-17D3Δ1 mutant animal. See also Figure 5-figure supplement 1. The movie was generated from heat-mapped time-series projection images of GCaMP6m fluorescence using MATLAB. Time and probe temperature are indicated at the top left and top right corners, respectively.

**Figure 1-source data 1.**

Source data used to generate the summary data and graphs shown in Figure 1 and Figure 1-figure supplement 1.

**Figure 4-source data 1.**

Source data used to generate the summary data and graphs shown in Figure 4 and Figure 4-figure supplement 1.

**Figure 5-source data 1.**

Source data used to generate the summary data and graphs shown in Figure 5 and Figure 5-figure supplement 1.

**Figure 7-source data 1.**

Source data used to generate the summary data and graphs shown in Figure 7.

**Figure 9-source data 1.**

Source data used to generate the summary data and graphs shown in Figure 9.

## Materials and Methods

### Fly strains

*y^1^w^1118^* strain was used as the control strain for *dsk*, *CCKLR-17D1*, and *CCKLR-17D3* mutants. *yv*; attp2 strain was used for the control of *yv; JF02644* (CCKLR-17D1 RNAi) and *yv; JF02968* (CCKLR-17D3 RNAi). *yw; nos-Cas9/CyO* (NIG-FLY CAS-0011) and *yw; Pr Dr/TM6C Sb Tb* were used for CRISPR/Cas9 mutagenesis. *Dp(3;1)2-2, w^1118^; Df(3R)2-2/TM3 Sb* (Bloomington #3688), *UAS-CCKLR-17D1* (66), and *UAS-CCKLR-17D3* (this study) were used for rescue experiments. *DSK-GAL4* (Bloomington #51981), *DSK-2A-GAL4* (Bloomington #84630), *R69E06-GAL4* (Bloomington #39493), *R69E06-lexA* (Bloomington #54925), *R72F11-lexA* (53), *R70F01-lexA* (Bloomington #53628), *R38A10-lexA* (Bloomington #54106), *R82A10-lexA* (Bloomington #54417), *CCKLR-17D1-T2A-GAL4* (79), *CCKLR-17D3-T2A-GAL4* (79) and *tsh-GAL80* were used for tissue-specific gene expressions. *40xUAS-IVS-mCD8::GFP* (Bloomington #32195), *10xUAS-IVS-mCD8GFP* (Bloomington #32186), *UAS>CD2 stop>mCD8::GFP hs-flp* (80), *lexA-rCD2::RFP UAS-mCD8::GFP* (81), *ppk-CD4::tdGFP* (Bloomington #35842), *UAS-nSyb-GFP_1-10_; lexAop-CD4-GFP_11_* (Bloomington #64314), *UAS-CD4-GFP_1-10_; lexAop-CD4-GFP_11_* (Bloomington #58755), *UAS-Denmark UAS-syt::eGFP* (Bloomington #33065), and *UAS-myrGFP QUAS-mtdTomato::3xHA; trans-Tango* (Bloomington #77124) were used for cellular visualizations. *UAS-GCaMP6m* (Bloomington #42748) was used for GCaMP Ca^2+^ imaging. *UAS-dTRPA1* (Bloomington #26263) (64) was used for thermogenetic experiments. *, UAS-CaMPARI2* (Bloomington #78316) and *lexAop-CaMPARI2* (Bloomington #78325) were used for CaMPARI2 snapshot imaging of neuronal activity. *dsk^sk1^*, *CCKLR-17D1* deletion mutants, *CCKLR-17D3* deletion mutants, and *UAS-CCKLR-17D3* were generated in this study and described below. *dsk*^attp^ strain (52) was obtained from the Bloomington stock center (#84497). Stocks were kept at 25°C with 12:12 hour light cycle on a standard food.

### Generating mutants and transgenic lines

Deletion mutants of *dsk*, *CCKLR-17D1,* and *CCKLR-17D3* were generated as described previously (82). Briefly, 23 bp-guide RNA (gRNA) sequences specific to the aimed region of the targeted genes were identified using an online tool (http://www.flyrnai.org/crispr/) and 20-bp sequences excluding PAM were cloned to the pBFv-U6.2 vector. Injections of the gRNA-pBFv-U6.2 vectors to yield transgenic fly strains were performed by BestGene Inc. The gRNA-expressing lines were crossed with a nos-Cas9 strain and about 20 independent F1 generations that could potentially possess modifications in the targeted genomic region were established. Lines that had a frameshifting deletion in the targeted region were screened through standard PCR and sequencing. The following gRNA and PCR primer sequences were used to generate *dsk*, *CCKLR-17D1*, and *CCKLR-17D3* mutants:

- *dsk*: GTAGACTAGTCGTCTGCGCT (gRNA), CCTCTAAACACTTGACAGCCGCGGTAACGG (forward primer), and CCGAAACGCATGTGACCGTAGTCATCG (reverse primer).
- *CCKLR-17D1*: GCTTCCGTGATACGCAGACTGGG (gRNA), ATGTGTTTTGTGGATACCCTGT (forward primer), and GGGCTATACCTCCA-TCAGTTTC (reverse primer).
- *CCKLR-17D3*: GCCATATCGGACATGCTGCTGGG (gRNA), GATAGGGA-TGGCTATATGGACACCGAGC (forward primer), and CTTAGCTGTCCCAATTCCCCCATCTTCT (reverse primer).

The UAS-CCKLR-17D3 line was generated through a ΦC31 integrase–based method. The oligo DNA corresponding to the sequence of CCKLR-17D3 mRNA (+447-2201 of AY231149) was synthesized (Eurofins Genomics) and cloned into the pUASg.attB vector (83) using the pENTR/D-TOPO Gateway cloning kit (Thermo Fisher Scientific, MA). The sequence-verified UAS-CCKLR-17D3 construct was integrated into the attP40 site to yield transgenic lines. The injections were performed by BestGene Inc.

### Thermal nociception assay

The experimenters were blinded to genotypes. Larval thermal nociception assays were performed as described previously (31), with slight modifications. A custom-made probe with a thermal feedback system was used. Unless otherwise noted, a thermal probe heated to 42°C was used to detect hypersensitive phenotypes (32). The results from all non-responders, defined as individuals that did not exhibit rolling behavior within 10 seconds, were converted to data points of 11 seconds for subsequent statistical analyses.

### Thermogenetic activation experiments

The experimenters were blinded to genotypes. A 60 mm dish containing approximately 1 ml distilled water (testing chamber) was placed on a temperature-controlled plate MATS-SPE (TOKAI HIT, Shizuoka, Japan) set at 44.5 °C to equilibrate the water temperature in a testing chamber to 35 ± 1 °C, which was continuously monitored using a T-type thermocouple wire IT-23 (Physitemp, NJ), USB-TC01 (National Instruments), and the NI Signal Express software (National Instruments, TX). Wandering third instar larvae expressing dTRPA1 by *ppk1.9-GAL4* and/or *DSK-2A-GAL4* were harvested to another 60 mm dish at room temperature (23-25 °C), and gently transferred to the 35 °C testing chamber using a paintbrush. All experiments were performed and recorded under a binocular microscope with a camcorder, and the latency from the placement of larvae to rolling was measured offline for each larva.

### Immunohistochemistry

The following antibodies were used in this study: chicken anti-GFP (Abcam, 1:500), mouse anti-GRASP (Sigma #G6539, 1:100) (62), mouse anti-rat CD2 (Bio-Rad, 1:200), rat anti-mCD8 (Caltag, 1:100), rabbit anti-FLRFa (a gift from Dr. Eve Marder, 1:5000) (84), rabbit anti-CD4 (Novus Biologicals, 1:500), mouse anti-REPO (Developmental Studies Hybridoma Bank 8D12, 1:5), rabbit anti-DsRed (Clontech #632496, 1:200), rat anti-HA (Roche 3F10, 1:100), mouse anti-CaMPARI-red (Absolute antibody 4F6, 1:1000), goat anti-HRP-Cy3 (Jackson ImmunoResearch, 1:100), goat anti-rat Alexa488 (Invitrogen, 1:500), goat anti-chicken Alexa488 (Invitrogen, 1:500), goat anti-mouse Alexa546 (Invitrogen, 1:500), goat anti-rabbit Alexa546 (Invitrogen, 1:500), goat anti-rat Alexa633 (Invitrogen, 1:500), and goat anti-rabbit Alexa633 (Invitrogen, 1:500). Dissected larval tissues were fixed in 4% paraformaldehyde for 30 minutes and then stained as previously described (32). The images were acquired by using a Zeiss LSM 510 with a 20x/0.75 Plan-Apochromat objective or 40x/1.0 Plan-Apochromat oil immersion objective, Zeiss LSM 700 with a 20x/0.75 Plan-Apochromat objective or 40x/1.0 Plan-Apochromat oil immersion objective, or Leica SP5 with a 40x/0.85 PL APO objective, and digitally processed using Zeiss LSM Image Browser, Leica LAS X Lite, and Adobe Photoshop.

### GRASP and *trans*-Tango experiments

The CD4-GRASP experiments were performed by crossing the *R69E06-lexA DSK-GAL4* strain with the *UAS-CD4-GFP_1-10_; lexAop-CD4-GFP_11_*. The nSyb-GRASP experiments were performed by crossing the *R69E06-lexA DSK-GAL4* with the *UAS-nSyb-GFP_1-10_; lexAop-CD4-GFP_11_*. Wandering 3rd instar larvae were dissected and immunostained as mentioned above. The signals of the reconstituted GFP were detected using a GFP antibody without cross-reaction to split-GFP fragments (Sigma #G6539) (62).

The *trans*-Tango experiments were performed by crossing the *R69E06-lexA, lexAop-rCD2::RFP UAS-mCD8::GFP; DSK-2A-GAL4* with *UAS-myrGFP QUAS-mtdTomato::3xHA; trans-Tango*. Wandering 3rd instar larvae were dissected and immunostained as mentioned above. The following combinations of the primary and secondary antibodies were used to avoid cross-contamination of fluorescent signals in microscopy: chick anti-GFP and anti-chick Alexa488, mouse anti-rCD2 and anti-mouse Alexa546, and rat anti-HA and anti-rat Alexa633.

### FLP-out clone analysis

The *UAS>CD2 stop>mCD8::GFP hs-flp* strain was crossed with *DSK-GAL4* or *DSK-2A-GAL4* to seed vials. Heat-shock induction of FLP-out clones and immunostaining were performed as described previously (32). The images of brain samples were acquired and digitally processed as described above.

### GCaMP calcium imaging

Ca^2+^ imaging of Goro neurons using GCaMP6m (85) was performed as described previously (32), with some modifications. Wandering third instar larvae expressing GCaMP6m in Goro neurons by *R69E06-GAL4* were dissected in ice-cold hemolymph-like saline 3.1 (HL3.1) (86) and imaged in a custom-made imaging chamber containing the HL3.1 equilibrated to the room temperature (23-25 °C). A Leica SP8 confocal microscope with resonant scanning system was used to perform three-dimensional time-lapse imaging. Z-stacks consisting of 4 to 6 optical slices of 512 x 512 pixel images were acquired at approximately 0.5 to 1 Hz using a 10x/0.4 PL APO objective lens with a zoom factor of 8.0. During imaging, a local heat ramp stimulation was applied to the lateral side of the A5 to A7 segment with a custom-made thermal probe. The probe temperature was controlled using a Variac transformer set at 20 V, which generated an approximately 0.6 °C/sec heat ramp stimulus. A T-type thermocouple wire was placed inside of the thermal probe to acquire the probe temperature readings and the data were acquired at 4 Hz through a digitizer USB-TC01 (National Instruments) and the NI Signal Express software (National Instruments). To minimize potential biases caused by day-to-day variations in imaging conditions and achieve fair comparisons, similar numbers of control and experimental genotypes were imaged side-by-side on the same day, using identical microscope settings. To monitor the temperature around the larval CNS during the local heat ramp stimulation, a small thermocouple wire (IT-23, Physitemp) was placed nearby the exposed CNS of semi-dissected larvae.

Maximum intensity projections were generated from the time-series Z-stacks on Leica LAS X Lite and the subsequent analyses of the images and temperature log data were performed using a custom-made code in MATLAB (MathWorks, MA). The region of interest (ROI) was selected as a circular area with a diameter of 15 pixels that covered the neurites of Goro neurons. Cell bodies were not used for the quantification because of small and variable increases in GCaMP6m signals upon heat stimulation. Average fluorescent intensity (F) was calculated for the ROI for each time point. The average of Fs from the first five frames was used as the baseline fluorescent intensity (F_0_) and percent changes of fluorescent intensity from the baseline [ΔF/F_0_ = (F - F_0_)*100/F_0_] was calculated for each time point. At least three ROIs were selected from areas that were expected to yield high GCaMP6m signal increases and the ROI that led to the highest peak ΔF/F_0_ was chosen for the subsequent statistical analysis. Samples whose highest peak ΔF/F_0_ was less than 10% were excluded from the analysis to avoid potential skewness of the data by inclusion of dead/unhealthy samples. Because the images and probe temperatures could not be acquired at the same time, probe temperature for each time point was calculated by linear interpolation from the raw readings. For comparisons among strains, ΔF/F_0_ data were binned and averaged in 1 °C intervals.

### CaMPARI2 experiments

Wandering third instar larvae expressing CaMPARI2 with *R69E06-GAL4* or *DSK2A-GAL4* were harvested from vials to a 60-mm petri dish and rinsed with water. Each larva was transferred onto kimwipe to remove moisture and attached with its ventral side up on labeling tape. A custom-made thermal probe (the same probe described for GCaMP experiments above) was placed at the lateral side of the A5 to A7 segment, and a local thermal ramp stimulus from room temperature (23-25 °C) to 50 °C was applied using a Variac transformer set at 20 V. Simultaneously with local heat ramp stimulation, 405 nm UV light was applied from 3 cm above the animal using a 405nm LED stand light (7 mW/cm^2^, LED405-9VIS, OPTOCODE Corporation, Tokyo, Japan). In “no heat” control experiments, no voltage was applied to the thermal probe attached to the larva, and only 405 nm light irradiation was applied for 40 seconds (equivalent to the length of time required to apply a temperature ramp stimulus from room temperature to 50°C with 20 V). In thermogenetic activation experiments, larvae expressing CaMPARI2 under the control of *R69E06-lexA* and dTRPA1 under the control of either only *ppk1.9-GAL4* or both *ppk1.9-GAL4* and *DSK-2A-GAL4* were immobilized on labeling tape with their ventral side up. The larvae were then transferred onto a MATS-SPE temperature-controlled plate set at 35 °C, and simultaneously irradiated with 405 nm light for 10 seconds from 3 cm above.

Immediately after thermal stimulation and/or photoconversion, the larvae were gently rinsed with water, removed from the labeling tape, and pooled into a new 60-mm dish. Five to fifteen animals were pooled for each experimental group and dissected immediately in ice-cold PBS. The larval CNS samples were immunostained following the protocol described above with mouse anti-CaMPARI red (4F6), rat anti-HA (3F10), anti-mouse Alexa 546, and anti-rat Alexa 633. The images of neurons expressing CaMPARI2 were acquired by using a Zeiss LSM 700 confocal microscope with a 40x/1.0 Plan-Apochromat oil immersion objective. All parameter settings of the microscope were kept identical across the experimental groups. To enhance green CaMPARI2 fluorescence, a 5% 405 nm laser was mixed with a 20% 639 nm laser. To achieve fair comparisons by minimizing the potential biases from day-to-day variations in stimulation, immunostaining, and imaging, similar numbers of animals from each experimental group were tested, immunostained, and imaged side-by-side on the same day.

Maximum intensity projections of images were generated from the Z-stacks using Zeiss Zen Lite software. Photoconversion rates of CaMPARI2 were quantified using MATLAB. Cell bodies of neurons expressing CaMPARI2 were automatically segmented using Otsu’s thresholding algorithm (87) based on the anti-HA signal intensity on the blue channel, and each image was cropped around the cell bodies with a margin of 50 pixels to the top, bottom, left, and right. Average intensities of green (green CaMPARI2) and red (anti-CaMPARI red) signals were calculated for cell bodies and the background (cropped image area excluding cell bodies), and the Red/Green ratio was calculated as follows: Red/Green ratio = (Average red signal intensity of cell bodies – Average red signal intensity of the background) / (Average green signal intensity of cell bodies – Average green signal intensity of the background). In the analysis of Goro neurons, images in which both sides of the cell bodies were segmented by Otsu’s method were included for the subsequent analyses.

### Data collection and statistical analyses

Mann–Whitney’s U-test was used for pair-wise comparisons and Steel’s test (the non-parametric equivalent of Dunnet’s test) or Steel–Dwass test (the non-parametric equivalent of Tukey-Kramer test) were used for multiple comparisons. Statistical analyses were performed in KyPlot 5.0 and GraphPad Prism 9. The numbers of samples (n) for all experiments indicate the numbers of biological replicates. Each experiment was repeated at least twice on different days to check the reproducibility, and all data were pooled for statistical analyses unless otherwise noted.

