## Supplemental figures for "A descending inhibitory mechanism of nociception mediated by an evolutionarily conserved neuropeptide system in *Drosophila*"

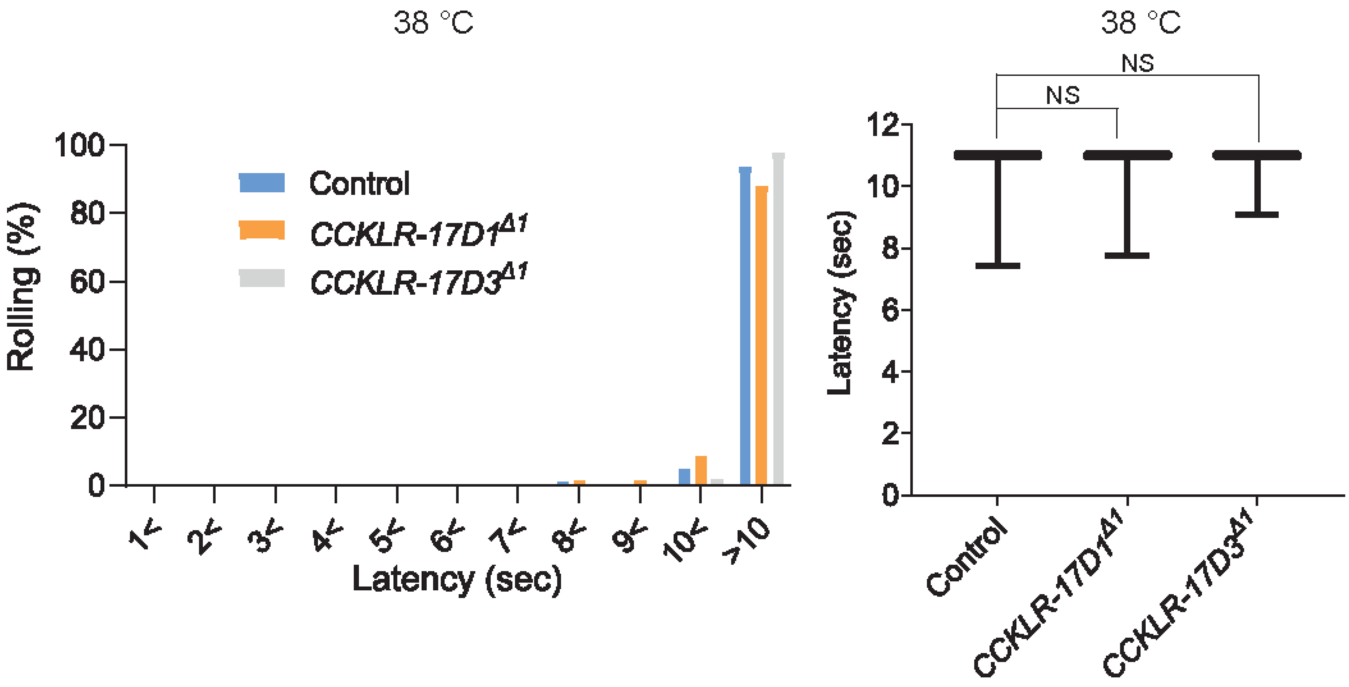


**Figure 1-figure supplement 1. Thermal nociceptive thresholds in DSK receptor mutants are largely normal.**

Thermal responses of *CCKLR-17D1* and *CCKLR-17D3* mutants to a 38 °C probe. Both *CCKLR-17D1*$^{\Delta 1}$ (n = 58) and *CCKLR-17D3*$^{\Delta 1}$ (n = 48) showed comparable distributions of responding latencies to the controls (*yw*, n = 79). Box plots show median (middle line) and 25th to 75th percentiles with whiskers indicating the smallest to the largest data points. $p > 0.4$ Steel's test.

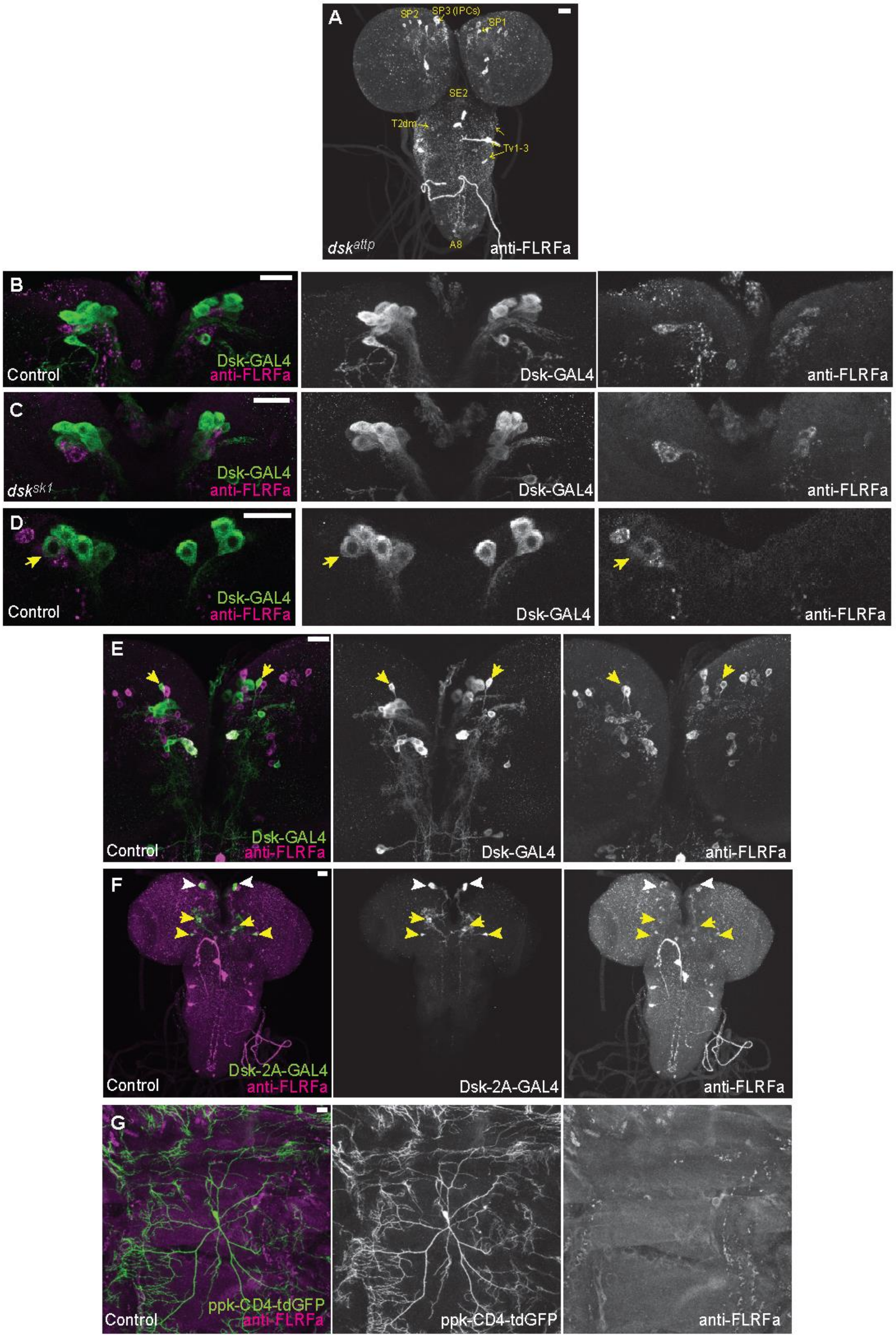
A
SP2
SP3 (IPCs)
SP1
SE2
T2dm
Tv1-3
A8
dsk^attp
anti-FLRFa
B
Control
Dsk-GAL4
anti-FLRFa
Dsk-GAL4
anti-FLRFa
C
dsk^sk1
Dsk-GAL4
anti-FLRFa
Dsk-GAL4
anti-FLRFa
D
Control
Dsk-GAL4
anti-FLRFa
Dsk-GAL4
anti-FLRFa
E
Control
Dsk-GAL4
anti-FLRFa
Dsk-GAL4
anti-FLRFa
F
Control
Dsk-2A-GAL4
anti-FLRFa
Dsk-2A-GAL4
anti-FLRFa
G
Control
ppk-CD4-tdGFP
anti-FLRFa
ppk-CD4-tdGFP
anti-FLRFa

7

**Figure 2-figure supplement 1. DSK expressions in the other brain neurons and peripheral sensory neurons.** (A) Representative image showing staining of anti-FLRFa in *dsk*$^{attp}$ mutants. The staining patterns were indistinguishable from *dsk*$^{sk1}$ mutants. (B) Representative image showing staining patterns of *DSK-GAL4* and anti-FLRFa in the wild-type larval IPCs. In the wild-type brain, typically 4 to 7 IPCs are visualized by *DSK-GAL4* and 1 to 3 by anti-FLRFa consistently to the previous study by Soderberg et al. [49] that used anti-DSK raised by Nichols et al. [47] In contrast, few IPCs co-labelled were found, which is also somewhat consistent with the variability of DSK immunostaining in IPCs reported by the previous study [49]. (C) Representative image showing staining patterns of *DSK-GAL4* and anti-FLRFa in the *dsk*$^{sk1}$ larval IPCs. Comparable staining patterns of *DSK-GAL4* and anti-FLRFa were observed in the mutant IPCs. (D) An example of rare IPC that was co-labeled by *DSK-GAL4* and anti-FLRFa in the wild-type brain (arrow). (E) Representative image showing SP1 neurons (arrows) that were rarely co-labeled by *DSK-GAL4* and anti-FLRFa. Note that faint anti-FLRFa staining was only detected in the SP1 neuron in the right hemisphere. (F) An example image showing *DSK-2A-GAL4* expression in IPCs (white arrowheads), MP1 (yellow arrows), and Sv neurons (yellow arrowheads). (G) Representative image showing double-labeling of Class IV md neurons (*ppk-CD4::tdGFP*) and anti-FLRFa staining in the dorsal larval body wall. All scale bars represent 20 μm.

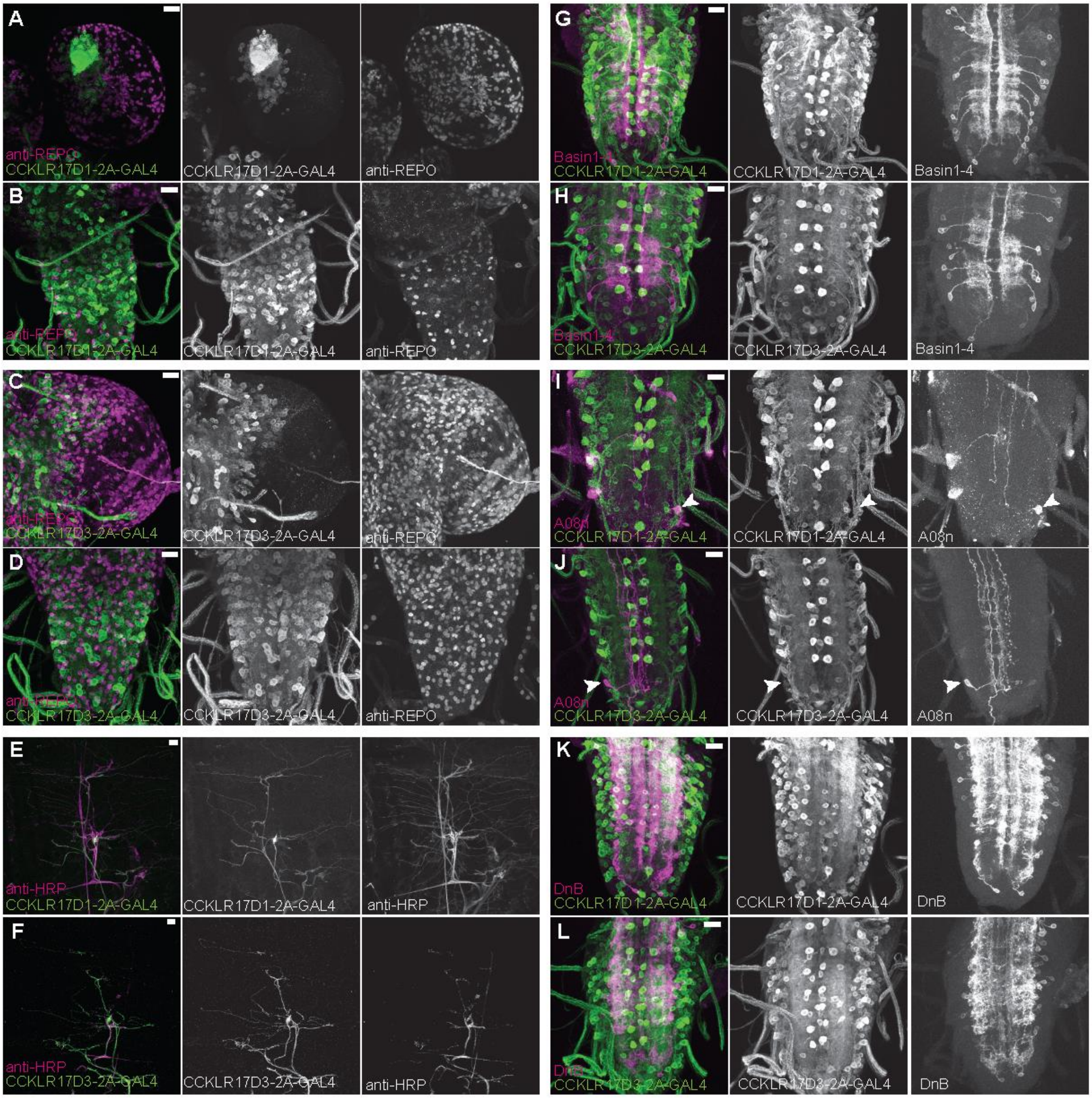


**Figure 3-figure supplement 1. Expression patterns of DSK receptors in larval glial cells, peripheral tissue, and nociceptive interneurons.**

(A and B) Representative images showing double-labeling of *CCKLR-17D1-T2A-GAL4* and a glial cell marker anti-

REPO in the larval brain (A) and the larval VNC (B), representing little expressions of *CCKLR-17D1-T2A-GAL4* in glial cells. (C and D) Representative images showing double-labeling of *CCKLR-17D3-T2A-GAL4* and a glial cell marker anti-REPO in the larval brain (C) and in the larval VNC (D), representing little expressions of *CCKLR-17D3-T2A-GAL4* in glial cells. (E and F) Representative images showing the expressions of *CCKLR-17D1-T2A-GAL4* (E) and *CCKLR-17D3-T2A-GAL4* (F) with a peripheral neuronal marker anti-HRP in the larval body wall, showing the absence of both GAL4s from peripheral sensory neurons other than es cells. (G and H) Expressions of *CCKLR-17D1-T2A-GAL4* (G) and *CCKLR-17D3-T2A-GAL4* (H) were negligible in Basin1 to 4 neurons labeled by *R72F11-lexA* in the larval VNC. (I and J) Negligible expression of *CCKLR-17D1-T2A-GAL4* (I) and *CCKLR-17D3-T2A-GAL4* (J) found in A08n neurons labeled by *R82E12-lexA* (arrowheads) in the larval VNC. (K and L) Expressions of *CCKLR-17D1-T2A-GAL4* (K) and *CCKLR-17D3-T2A-GAL4* (L) were negligible in DnB neurons labeled by *R70F01-lexA* in the larval VNC. All scale bars represent 20 µm.

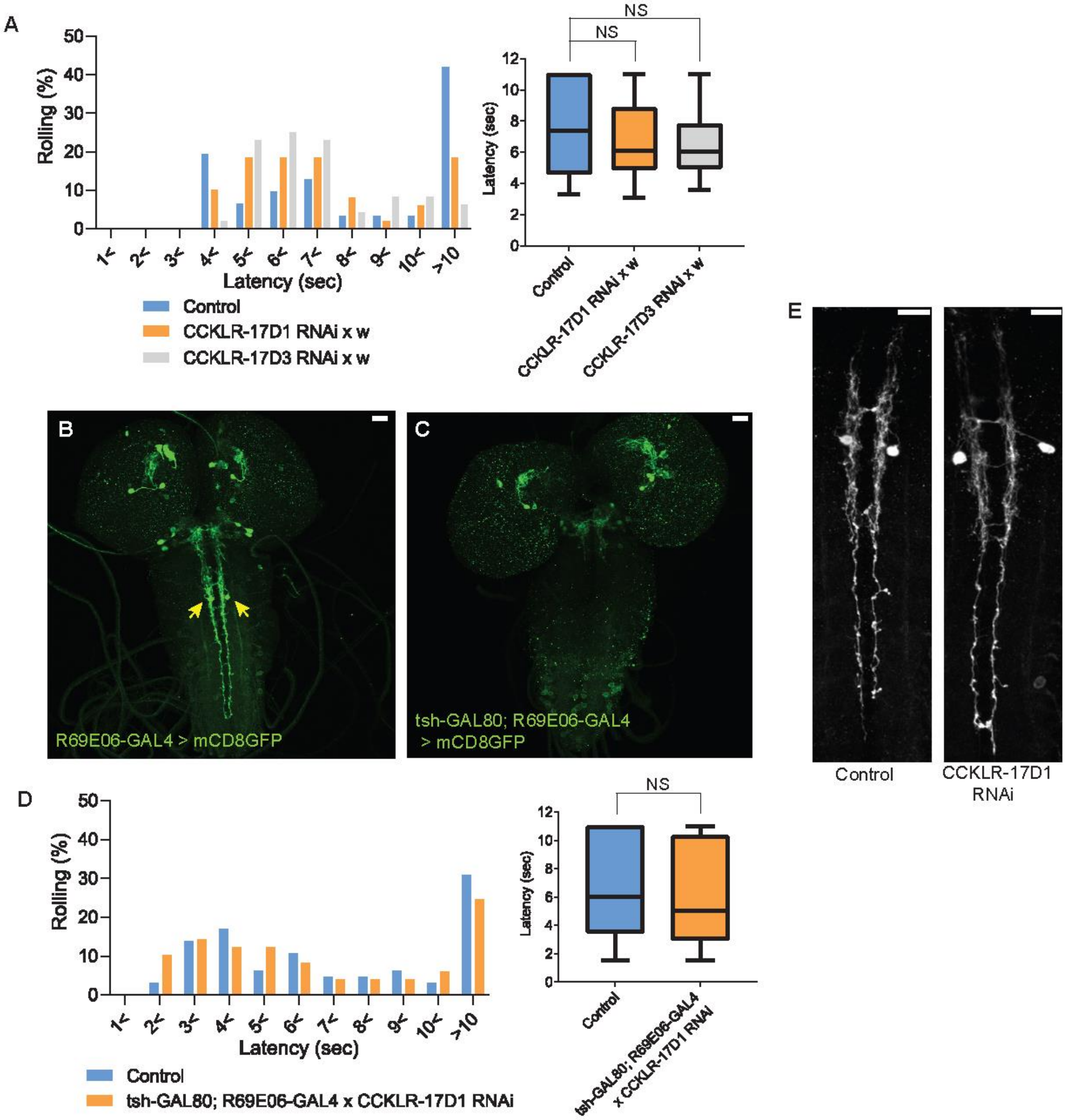


**Figure 4-figure supplement 1. CCKLR-17D1 RNAi in Goro neurons induces thermal hypersensitivity in a GAL4-dependent manner, without affecting the morphology.**

(A) Without the GAL4 driver, UAS-CCKLR-17D1 RNAi (*yv;JF02644* x $w^{1118}$, n = 49) and UAS-CCKLR-17D3 RNAi (*yv;JF02968* x $w^{1118}$, n = 48) both did not cause thermal hypersensitivity to 42 °C compared with the control (*R69E06-GAL4* x *yv; attp2*, n = 31). $p > 0.38$ Steel's test. (B and C) *tsh-GAL80* eliminates the expression of *R69E06-*

*GAL4* from Goro neurons. The expression of *R69E06-GAL4* in Goro neurons (B) (*10xUAS-IVS-mCD8GFP; R69E06-GAL4* x *yv; attp2*) was eliminated when *tsh-GAL80* was combined with *R69E06-GAL4* (C) (*tsh-GAL80; R69E06-GAL4* x *40xUAS-IVS-mCD8GFP*). (D) When the expression of UAS-CCKLR-17D1 RNAi was eliminated from Goro neurons (*tsh-GAL80; R69E06-GAL4* x *yv; JF02644*, n = 49), larval nociceptive responses to a 42 °C probe was indistinguishable from the control (*R69E06-GAL4* x *yv; attp2*, n = 65). $p > 0.33$ Mann-Whitney's U-test. (E) Goro neurons expressing CCKLR-17D1 RNAi (*10xUAS-IVS-mCD8GFP; R69E06-GAL4* x *yv; JF02644*) did not alter their number, position, and gross projection patterns in comparison with the control (*10xUAS-IVS-mCD8GFP; R69E06-GAL4* x *yv; attp2*). All scale bars represent 20 µm. All box plots show median (middle line) and 25th to 75th percentiles with whiskers indicating the smallest to the largest data points.

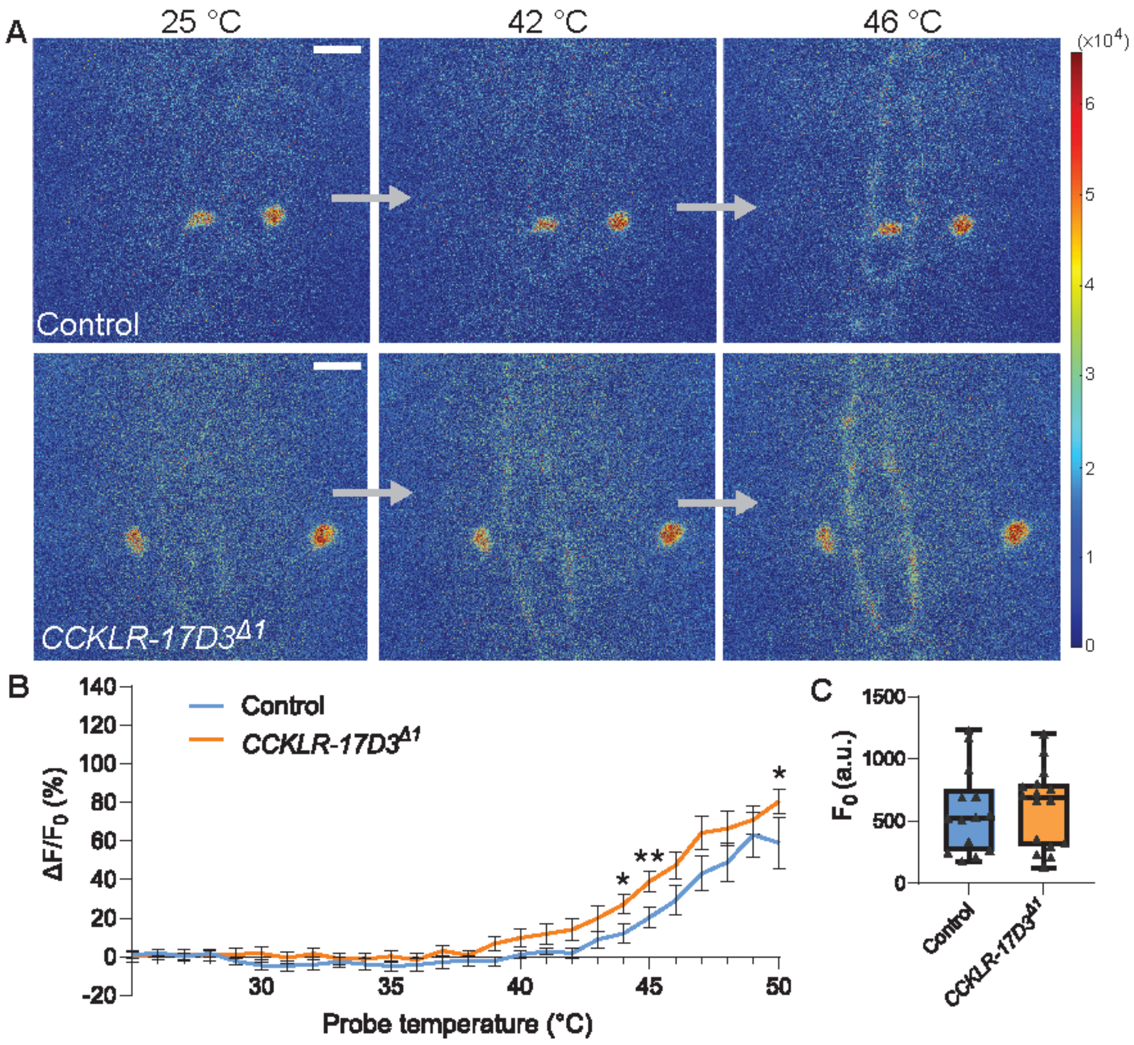


**Figure 5-figure supplement 1. Thermal responsiveness of Goro neurons in CCKLR-17D3Δ1.**

(A) Representative still images showing thermal activation of Goro neurons in control animals (top, *yw*/Y; *R69E06-GAL4 UAS-GCaMP6m*/+) and *CCKLR-17D3*$^{\Delta 1}$ mutants (bottom, *CCKLR-17D3*$^{\Delta 1}$*/Y; R69E06-GAL4 UAS-GCaMP6m*/+). See also Movie S5 and S6. Scale bars represent 20 µm. (B) Average percent increase of GCaMP6m fluorescence intensity relative to baseline ($\Delta F/F_0$) during heat ramp stimulations in *CCKLR-17D3*$^{\Delta 1}$ experiments. $\Delta F/F_0$ is plotted to binned probe temperature (interval = 1 °C). In comparison with controls, Goro neurons of *CCKLR-17D3*$^{\Delta 1}$ exhibited mildly elevated fluorescent increase of GCaMP6m, which reached statistical significance only at 44°C, 45°C, and 50°C. n = 14 and 16 for controls and *CCKLR-17D3*$^{\Delta 1}$, respectively. * $p < 0.05$, ** $p < 0.01$ (Mann–Whitney's U-test). Error bars represent standard errors. (C) Basal GCaMP6m signal levels ($F_0$) did not differ between controls and *CCKLR-17D3*$^{\Delta 1}$ (n = 14 and 16). $p > 0.5$ (Mann–Whitney's U-test). Box plots show median (middle line) and 25th to 75th percentiles with whiskers indicating the smallest to the largest data points.
